# Temporally and regionally distinct morphogenetic processes govern zebrafish tail fin blood vessel network expansion

**DOI:** 10.1101/2022.06.14.496153

**Authors:** Elvin V. Leonard, Sana Safatul Hasan, Arndt F. Siekmann

## Abstract

Blood vessels form elaborate networks depending on tissue-specific signalling pathways and anatomical structures to guide their growth. However, it is not clear which morphogenetic principles organize the stepwise assembly of the vasculature. We thus performed a longitudinal analysis of zebrafish tail fin vascular assembly, revealing the existence of temporally and spatially distinct morphogenetic processes. Initially, vein-derived endothelial cells (ECs) generated arteries in a reiterative process requiring Vascular Endothelial Growth Factor (VEGF), Notch and *cxcr4a* signalling. Subsequently, veins produced veins in more proximal fin regions, transforming pre-existing artery-vein loops into a three-vessel pattern consisting of an artery and two veins. A distinct set of vascular plexuses formed at the base of the fin. They differed by virtue of diameter, flow magnitude and marker gene expression. At later stages, intussusceptive angiogenesis occurred from veins in distal fin regions. In proximal fin regions, we observed new vein sprouts crossing the inter-ray tissue through sprouting angiogenesis. Together, our results reveal a surprising diversity among the mechanisms generating the mature fin vasculature and suggest that these might be driven by separate local cues.

## Introduction

All organs and tissues need to be vascularized during embryonic development to ensure their proper function later in life. Work over the last decades has revealed a remarkable diversity in terms of anatomy and genetic make-up of organ-specific blood vessels, tailoring them to the needs of the organ they vascularize (Aird, 2007a; b; Augustin and Koh, 2017; Eelen et al., 2020). This is supported by single cell sequencing of mouse tissues revealing that EC heterogeneity can be more strongly correlated with different tissues instead of blood vessel types (Kalucka et al., 2020; Paik et al., 2020). Prominent examples of tissues containing ECs with particular properties are the brain (Langen et al., 2019) and the liver (Koch et al., 2021). Lymphatics also possess tissue-specific properties (Petrova and Koh, 2018). Species-specific vascular structures to meet habitat and life-style challenges also exist. Tuna fish evolved special vascular beds that coordinate heat exchange between a large set of arterioles and venules, necessary for maintaining optimal muscle temperature and swimming performance in colder waters (Stevens et al., 1974). Dedicated lymphatic structures can provide hydraulic control of fin shapes (Pavlov et al., 2017). In addition, proper organ development relies on the exchange of signalling molecules between ECs and their surrounding cells even before the onset of blood flow in a process referred to as angiocrine signalling (Gomez-Salinero et al., 2021; Rafii et al., 2016). For instance, in the central nervous system specialised ECs initially organize neurogenesis (Tata and Ruhrberg, 2018) followed by blood-brain barrier formation (Andreone et al., 2015). A similar crosstalk instructs bone formation (Sivan et al., 2019). Therefore, intricate interactions between the endothelium and other tissue cell types are instrumental for proper organogenesis and tissue function.

Despite these insights, our understanding of the morphogenetic principles that govern the formation of organ-and tissue-specific blood vessels over time and how they might be intertwined with organ function is limited. We also do not know whether regenerating tissues re-use developmental programs (Goldman and Poss, 2020). As a first step towards an understanding of these mechanisms, we decided to study the stepwise assembly of the zebrafish tail fin vasculature. Previous studies have shown that this vasculature forms in a stereotypical pattern, where each fin ray harbours a central artery that is located within the ray’s bone and is flanked by two lateral veins located on either side of the bone ray (Huang et al., 2003; Kametani et al., 2015; Paulissen et al., 2022; Xu et al., 2014). Imaging studies performed during fin regeneration elucidated that pre-existing veins give rise to regenerating arteries in a process that depends on *cxcr4a* (Xu *et al*., 2014) and Notch signalling (Kametani *et al*., 2015). Elegant lineage tracing studies conducted in regenerating fins similarly provided evidence that arterial and venous ECs share a common lineage precursor cell (Tu and Johnson, 2011). While a vascular plexus is present at the tips of regenerating fin vessels, this plexus is either absent or greatly reduced in normal growing fins (Huang et al., 2009; Xu *et al*., 2014), suggesting that distinct processes might pattern regenerating adult blood vessels in comparison to blood vessels in developing fish. The formation of this plexus further depends on structural features of the extracellular matrix, as interfering with collagen assembly results in plexus defects (Huang *et al*., 2003; Huang *et al*., 2009; Nakagawa et al., 2022; Senk and Djonov, 2021). A recent report showed that osteoblasts are required for normal fin blood vessel outgrowth (Bump et al., 2022). This indicates that osteoblasts likely secrete pro-angiogenic factors, as also proposed by Paulissen et al. (Paulissen *et al*., 2022). We previously showed that the pro-angiogenic chemokine *cxcl12a* is expressed in central fin regions and can accumulate in the bone (Xu et al., 2014). Thus, structural features of the fin can affect local blood vessel patterning.

Fin blood vessel formation requires Vascular Endothelial Growth Factor (VEGF) signalling (Bayliss et al., 2006; Bump *et al*., 2022). Fins lacking blood vessels did not show overt patterning defects in bone or nerve fibres but failed to grow after the initial stages of morphogenesis (Bump *et al*., 2022). This was more pronounced during fin regeneration (Bayliss *et al*., 2006). Therefore, it remains to be determined whether blood vessels instruct fin morphogenesis in a subtle manner, leading to later defects in growth or whether these defects are caused by a lack of nutrient and oxygen supply. Recent results further revealed surprising differences in the formation of blood vessels among different types of zebrafish fins. Das et al. showed that lymphatics are precursor cells for anal fin blood vessels (Das et al., 2022), while Paulissen et al. showed that the vasculature in pectoral fins was derived from pre-existing blood vessels (Paulissen *et al*., 2022).

To better understand the processes governing the step-by-step assembly of an organ-specific vascular bed, we set out to examine blood vessel formation throughout tail fin development. Our findings reveal that initial blood vessel growth occurs through a reiterative set of angiogenic sprouts emanating from veins and giving rise to veins and arteries in a VEGF and *cxcr4a*-dependent manner. A later occurring set of vein-derived sprouts exclusively generates veins, establishing a 3-vessel pattern. This is further elaborated through inter-vessel connections, which are formed through sprouting and intussusceptive angiogenesis. We also detected a separate vessel network that formed a plexus at the base of the fin and could be distinguished in terms of vessel diameters and transgene expression levels. This network formed though sprouting of blind-ended, lumenized vessels. Together, our results reveal a marked diversity in the morphogenetic principles occurring at different anatomical locations and developmental stages of fin growth while generating distinct fin blood vessel types. They also lay the groundwork for the further examination of their functional properties and the molecules controlling their formation.

## Results

### Successive waves of angiogenic sprouting generate the primitive fin vasculature

To investigate tail fin vascular morphogenesis in detail, we used *Tg(fli1a:nEGFP)*^*y7*^; *(−0*.*8flt1:RFP)*^*hu5333*^ double transgenic embryos. In these, all EC nuclei are labelled by virtue of EGFP expression, while arterial ECs more strongly express RFP. We examined vascular growth starting at 2.5 dpf (Fig. 1A). We observed a single blood vessel sprout, emanating from the posterior cardinal vein (PCV; Fig. 1B). Of note, at the distal end of the sprout, several ECs had switched on RFP expression, indicative of the onset of arterial specification (Fig. 1B, arrowheads). By 1 wpf, we detected the presence of a new circulatory loop of axial vessels extending posteriorly (Fig. 1C, bracket). Furthermore, a new blood vessel sprouted from this loop towards the ventral region of the growing fin fold (Fig. 1C, arrowhead). Additional branching was initiated from these sprouts, resulting in the extensive expansion of the vascular tree by 2 wpf (Fig. 1D).

**Figure 1.**
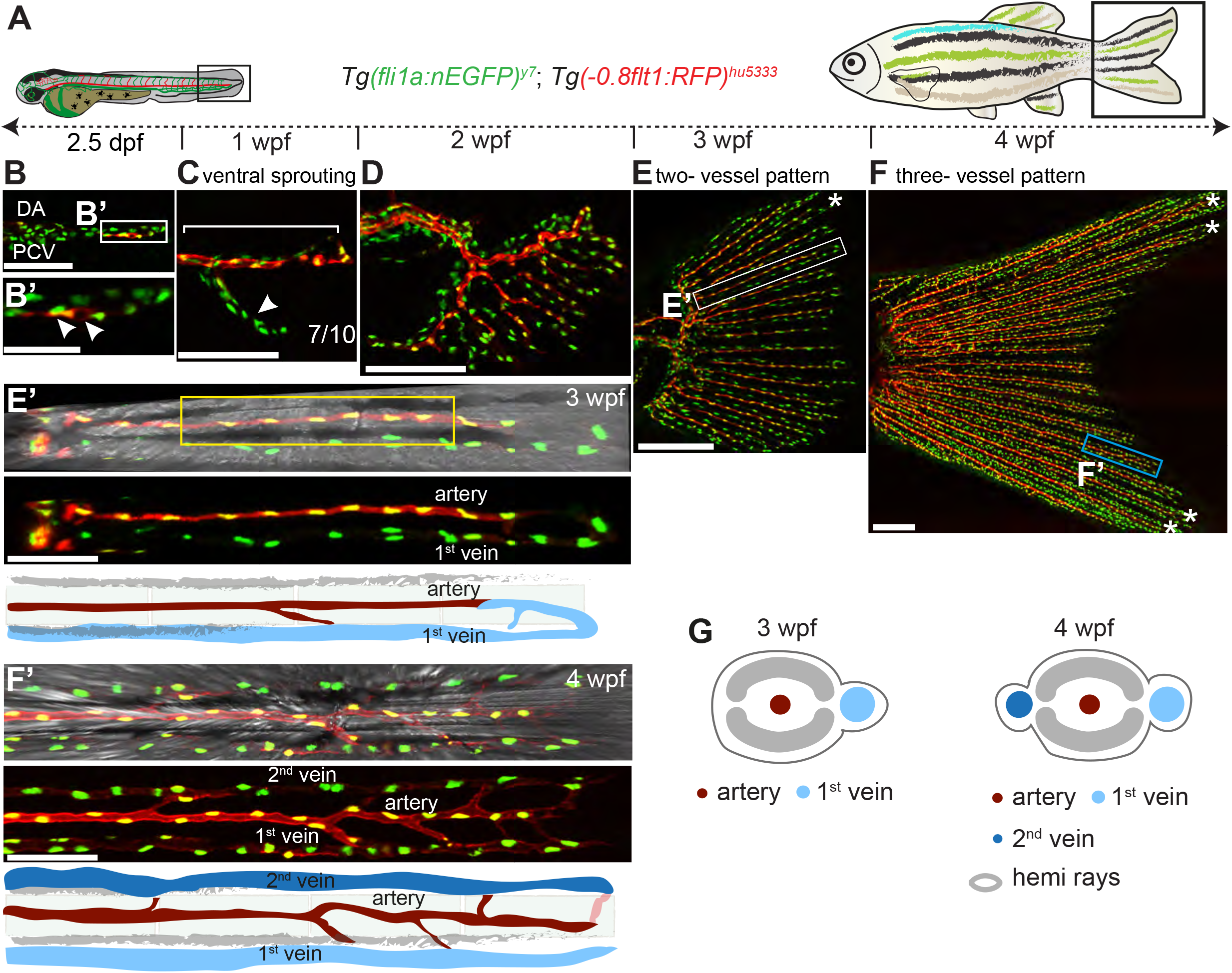
Branching morphogenesis of zebrafish caudal fin vasculature. **(A)** Schematic representation of the time course of the study. Caudal fin vascular development was studied from 2.5 dpf to 4 wpf. **(B)** Maximum intensity projections of confocal z-stacks of *Tg(fli1a:nEGFP)*^*y7*^; *Tg(−0*.*8flt1:RFP)*^*hu5333*^ double transgenic fish labelling all EC nuclei (green) and arteries (red). Initial sprouting from PCV to form posterior axial vessels at 2.5 dpf. Scale bar: 20 μm. Boxed region enlarged in (**B’**) indicates onset of arterial specification (white arrowheads). Scale bar: 5 μm. **(C)** First ventral sprout (white arrowhead) extending into the fin fold at 1 wpf. Bracket marks posterior axial vessels. Scale bar: 20 μm. **(D)** Further branching and expansion of the vascular tree into the caudal fin at 2 wpf. Scale bar: 20 μm. **(E)** Alignment of developing vasculature with fin rays at 3 wpf. Scale bar: 50 μm. Boxed area enlarged in **(E’)** shows two-vessel pattern (artery-vein), where the artery is aligned to the centre of the bone ray (yellow box) with the vein on one side. Top image includes bright field channel to visualize bone rays. Scale bar: 20 μm. **(F)** Caudal fin vasculature at 4 wpf. Scale bar: 100 μm. Boxed region enlarged in (**F’**) shows caudal fin vasculature displaying a three-vessel pattern (vein-artery-vein) with a medially located artery flanked by a vein on either side. Top image includes bright field channel to visualize bone rays. Scale bar: 20 μm. Numbers represent individual embryos (or fish) analysed for each developmental stage. **(G)** Schematic representation of two and three-vessel patterns.

Subsequently, these vessels organized into a radial pattern, where each fin ray was inundated by an artery and a vein, which came to lie next to each other, resulting in a two-vessel pattern (Fig. 1E,E’). Arteries were located within the forming bone rays, while veins were positioned outside of the bone rays. By 4 wpf, the vasculature had matured into a three-vessel structure, where a central artery was flanked by two veins (Fig. 1F,F’). While arteries were located within forming bone rays, veins were positioned outside of the bone rays (Fig. 1G). Thus, the vasculature of the fin matures from an initial loop, through which each artery drains into a single vein, into a configuration, where each artery drains into two veins (Figure 1G).

### Vein-derived angiogenic sprouts initially generate new arteries

To understand the mechanism through which the fin vasculature expands, we performed live imaging for 16 hours (h) starting at 52 hpf using *Tg(fli1a:nEGFP)*^*y7*^; *Tg(−0*.*8flt1:RFP)*^*hu5333*^ double transgenic embryos (Fig. 2A, Supplementary Fig. 1A,B, Supplementary Movies 1-3). We could distinguish individual ECs that initially sprouted from the PCV and migrated horizontally toward the tip of the tail (Cells 1 and 2, 0 h time point). Eventually, ECs at the leading edge of the sprout acquired arterial identity, as evidenced by onset of *Tg(−0*.*8flt1:RFP)*^*hu5333*^ expression (Fig. 2A, cell1 at 3 h time point and cells 1 and 2, 10 h time point, arrowheads). These ECs then migrated in the reverse direction along the existing vasculature until they reached the DA, where they anastomosed (Fig. 2A, 16 h time point). To corroborate our findings, we imaged the nascent fin vasculature at different time points in *Tg3(fli1a:lifeactGFP)*^*mu240*^; *Tg(−0*.*8flt1:RFP)*^*hu5333*^ double transgenic animals. As before, we observed the onset of RFP expression in ECs that had just started to turn in their migration direction and before connecting to the pre-existing artery (Supplementary Fig. 2A-D, red arrowheads). Subsequently, *flt1:RFP* expressing ECs connected to the tail artery, while new cells positioned at the distal sprout tip started to express *flt1:RFP* and turned around (Supplementary Fig. 2E,E’, red arrowheads). Thus, while the tail vein elongated in a proximal to distal manner, the tail artery extended through the continuous addition of previously distally located cells that initially were part of the vein sprout.

**Figure 2.**
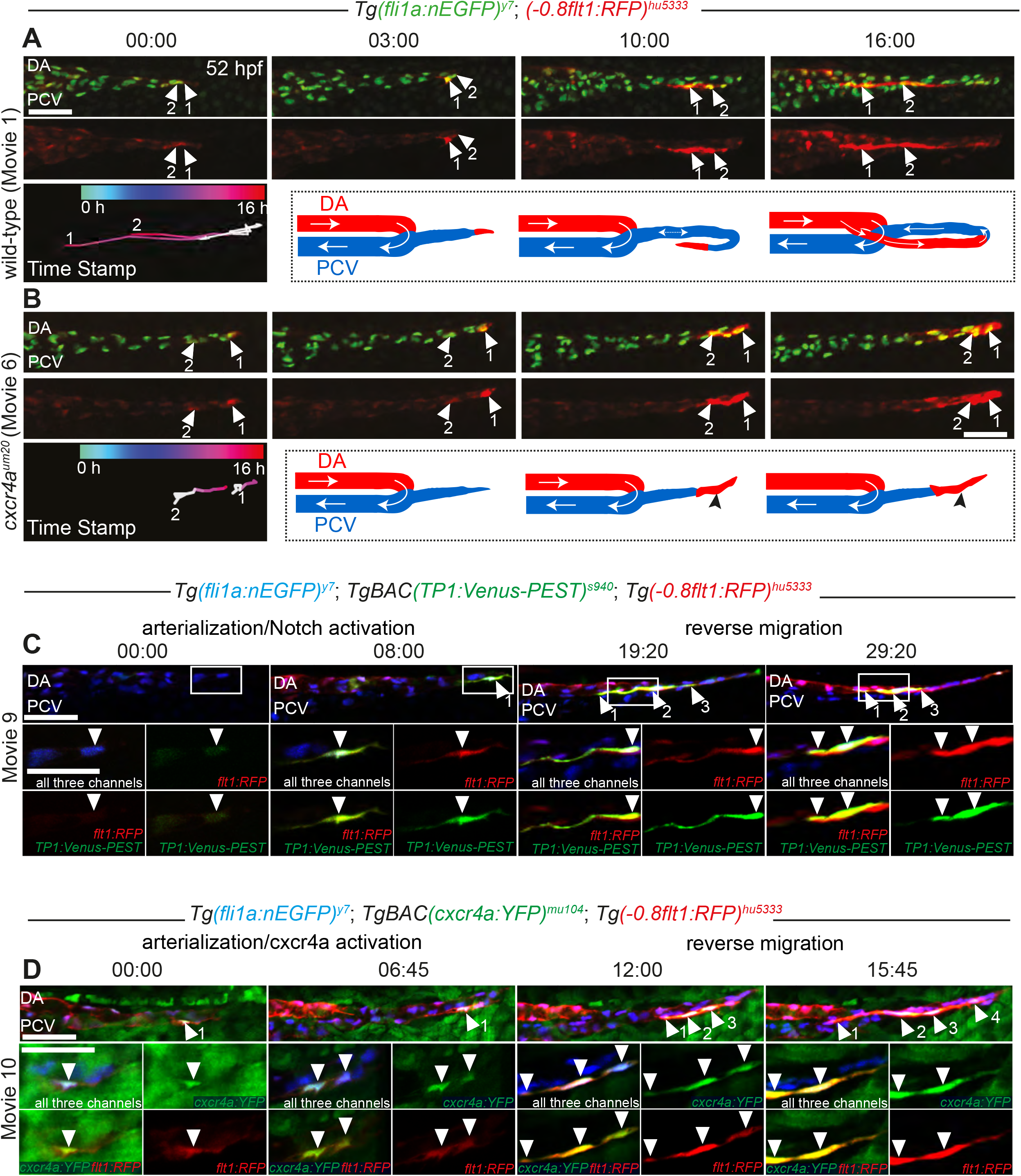
ECs within the first circulatory loop of the fin vasculature become arterialized and activate *cxcr4a* and Notch signalling. **(A)** Still images at indicated time points of ECs sprouting from PCV in *Tg(fli1a:nEGFP)*^*y7*^; *Tg(−0*.*8flt1:RFP)*^*hu5333*^ double transgenic wild-type control animal, labelling EC nuclei (green) and arterial ECs (red). White arrowheads with numbers mark individual ECs. Time encoded migration tracks of marked ECs. t0=52 hpf. Schematic representation of sprout formation, white arrows indicate direction of blood flow. Scale bar: 50 μm. **(B)** Still images at indicated time points of ECs sprouting from PCV in *Tg(fli1a:nEGFP)*^*y7*^; *Tg(−0*.*8flt1:RFP)*^*hu5333*^ double transgenic *cxcr4a*^*um20*^ mutants. Schematic representation of observed phenotype in *cxcr4a* mutants. Black arrowheads indicate clustering of arterial ECs at the tip of the sprout. Scale bar: 50 μm. **(C)** Maximum intensity projections of confocal z-stacks of *Tg(fli1a:nEGFP)*^*y7*^; *Tg(−0*.*8flt1:RFP)*^*hu5333*^; *Tg(TP1:Venus-Pest)*^*s940*^ triple transgenic fish labelling all EC nuclei (blue), arterial ECs (red) and cells with Notch pathway activation (green) in lateral views anterior to the left. White arrowheads indicate arterial ECs with activated Notch signalling. Scale bar: 20 μm. Boxed regions are enlarged. Scale bar: 4 μm. **(D)** Still images at indicated time points of ECs sprouting from PCV in *Tg(fli1a:nEGFP)*^*y7*^; *Tg(cxcr4a:YFP)*^*mu104*^; *Tg(−0*.*8flt1:RFP)*^*hu5333*^ triple transgenic embryos labelling all EC nuclei (blue), arterial ECs (red) and *cxcr4a* positive cells (green) in lateral views anterior to the left. White arrowheads indicate ECs that express *cxcr4a* and show onset of arterial specification. Scale bar: 20 μm. Boxed regions are enlarged. Scale bar: 4 μm.

We then used *Tg(gata1a:DsRed)*^*sd2*^ fish to visualize red blood cells and blood flow patterns. The nascent vein sprout was lumenized from very early stages onwards, while still exclusively being connected to the venous circulation, as supported by the presence of red blood cells within the sprout (Supplementary Fig. 2F-H, yellow arrowheads). As the sprout lacked an outlet, we observed red blood cells oscillating within the sprout (Supplementary Movies 4,5). The sprout eventually fused with the pre-existing artery, leading to the onset of blood flow through the newly connected artery and vein (Supplementary Fig. 2I-I’). Furthermore, after completion of the arterial connection, the flow pattern within the sprout changed from being oscillatory to laminar. Therefore, this morphogenetic process generates a figure-8 configuration of tail and trunk arteries and veins. In the fish trunk, the artery is in a dorsal position with respect to the trunk vein, while in the area later inundating the tail fin the artery runs ventrally in respect to the tail vein (Fig. 2I, schematic).

### Vegf and Notch signalling upstream of *cxcr4a*-dependent EC migration control fin artery morphogenesis

Of interest, these morphogenetic processes strikingly resembled those that we previously observed during formation of the eye vasculature in embryonic zebrafish (Hasan et al., 2017). Here we showed that Vegf and Notch signalling upstream of *cxcr4a* were important for guiding vein-derived arterial sprouting. We therefore investigated whether these signalling molecules similarly control fin vascular expansion. Time-lapse analysis of artery formation in *cxcr4a*^*um20*^ mutant zebrafish showed normal expression of the arterial marker *flt1:RFP*, but failure of these cells to migrate towards the more proximally located artery (Fig. 2B, Supplementary Fig. 1C,D, Supplementary Movies 6-8). Instead, we found arterial-fated cells accumulating at the distal tip of the newly sprouting tail fin sprout (Fig. 2B, compare 10 and 16 h time point to wild-type 10 and 16 h time point in Fig. 2A, compare also schematic drawings in Fig. 2B with schematic in Fig. 2A). Thus, like in the embryonic eye vasculature, *cxcr4a* signalling is instrumental in guiding the nascent artery sprout in the tail fin towards the dorsal aorta without affecting arterial specification.

To examine the onset of Notch pathway activation and *cxcr4a* expression within the growing sprout, we performed time-lapse imaging in triple transgenic animals. To monitor Notch pathway activation, we used the Notch indicator line *Tg(TP1:Venus-PEST)*^*s940*^ together with *Tg(−0*.*8flt1:RFP)*^*hu5333*^ and *Tg(fli1a:nEGFP)*^*y7*^. Time-lapse imaging revealed active Notch signalling in ECs that also expressed *flt1:RFP* at the tip of the reversing artery sprout (Fig. 2C, see also Supplementary Movie 9). We further confirmed *cxcr4a* expression in arterial ECs by performing time-lapse imaging of triple transgenic *Tg(−0*.*8flt1:RFP)*^*hu5333*^; *(cxcr4a:YFP)*^*mu104*^; *Tg(fli1a:nEGFP)*^*y7*^ animals. These movies revealed an overlap between *cxcr4a* and *flt1* expression (Fig. 2D, Supplementary Movie 10). We furthermore detected *cxcr4a* expression in ECs with activated Notch signalling (Supplementary Fig. 3A, Supplementary Movie 11). Thus, like ECs in other angiogenic settings, vein cells populating the tail fin in zebrafish larvae can differentiate into the arterial lineage, initiate Notch signalling and express *cxcr4a* (Fig. 3D).

**Figure 3.**
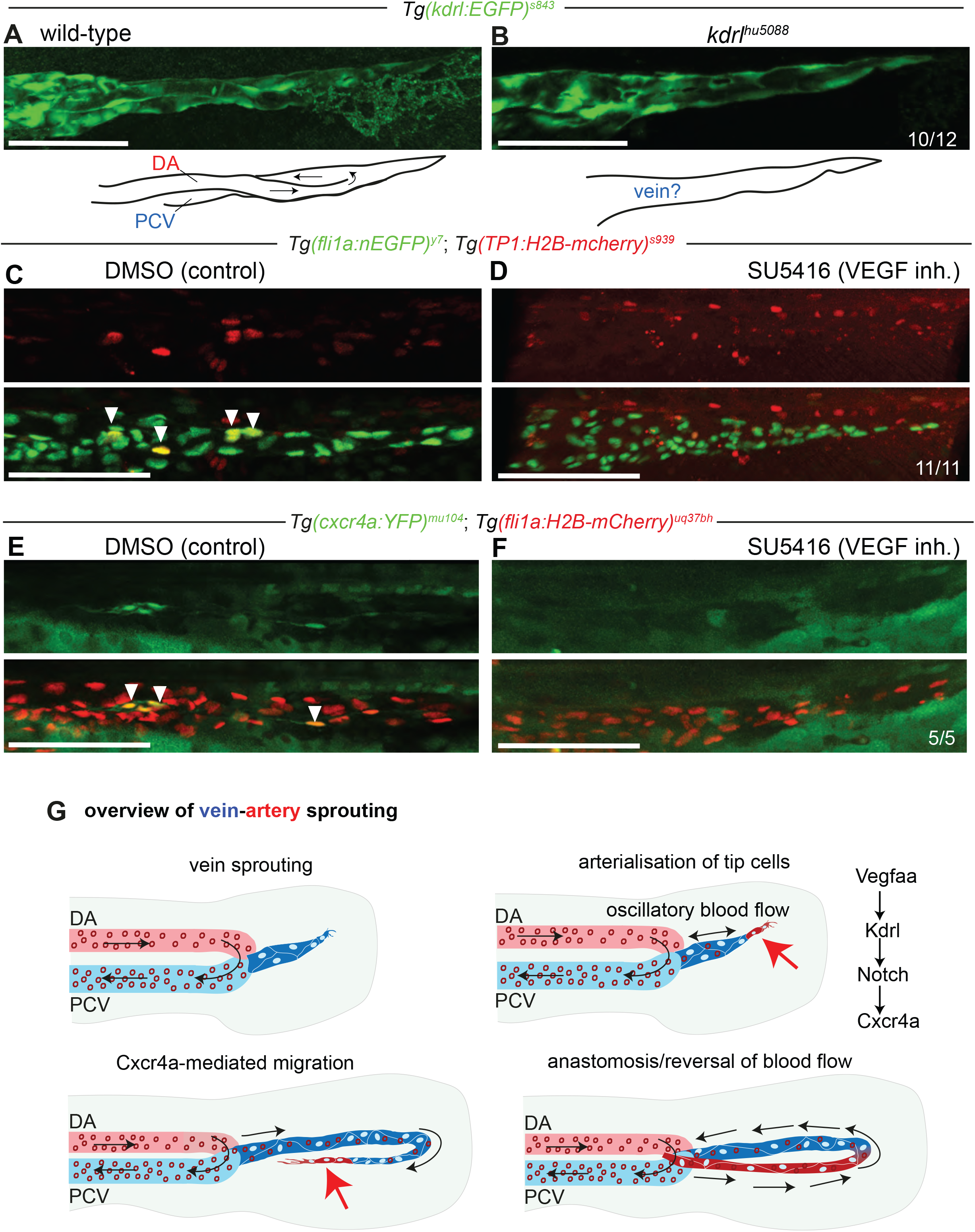
Sprouting of the first circulatory loop depends on VEGF signalling upstream of Notch signalling. **(A)** Maximum intensity projection of confocal z-stacks of *Tg(kdrl:EGFP)*^*s843*^, wild-type control. Scale bar: 20 μm. Schematic drawing indicates arteries and veins in tail blood vessel sprout. **(B)** Maximum intensity projection of confocal z-stacks of *Tg(kdrl:EGFP)*^*s843*^, *kdrl*^*hu5088*^ mutant. Schematic drawing indicates tail blood vessel sprout lacking discernible arteries. Scale bar: 15 μm. **(C)** Maximum intensity projections of confocal z-stacks of *Tg(fli1a:nEGFP)*^*y7*^; *Tg (TP1:H2B-mcherry)*^*s939*^, DMSO control treatment. White arrowheads indicate ECs with activated Notch pathway. **(D)** SU5416 treatment causes loss of Notch pathway activation in ECs. Scale bar: 10 μm. **(E)** Maximum intensity projection of confocal z-stacks of *Tg(cxcr4a:YFP)*^*mu104*^; *Tg (fli1a:H2B-mCherry)*^*uq37bh*^, DMSO control. White arrowheads indicate *cxcr4a* expression in ECs. **(F)** SU5416 treatment leads to loss of *cxcr4a* expression. Scale bar: 10 μm. **(G)** Schematic representation and overview of signalling pathways involved in controlling vein-artery sprouting in the tail fin.

To functionally test for the requirement of VEGF signalling during artery formation, we first analysed tail fin vascularization in zebrafish larvae with compromised Vegf signalling. Mutants for either the Vegfa receptor *kdrl* (Fig. 3A,B) or the Vegfaa ligand (Supplementary Figure 3B,C) displayed normal outgrowth of the vein sprout into the tail but lacked arterial cells that would display a turning behaviour. We made similar observations in larvae treated with the Vegf inhibitor SU5416 (Fig. 3C,D). We then used this inhibitor to test for Notch pathway activation downstream of Vegf. We observed absence of expression of the Notch pathway indicator line *Tg(TP1:H2B-mcherry)*^*s939*^ in inhibitor treated embryos (Fig. 3C,D), demonstrating that Notch pathway activation requires Vegf signalling in tail fin ECs. We also observed absence of *Tg(cxcr4a:YFP)*^*mu104*^ expression in SU5416 treated embryos (Fig. 3E,F). Together, these results indicate that the mechanisms guiding the formation of the initial artery-vein circulatory loop inundating the tail fin are like those operating in other regions of the embryo. We now term this mode of angiogenic sprouting “vein-artery sprouting” (Fig. 3G).

### Expansion of the fin vasculature occurs at two distinct anatomical regions

We followed the further expansion of the fin vasculature, which occurred in several distinct anatomical locations. In order to do so, we used *Tg(−0*.*8flt1:RFP)*^*hu5333*^ fish to label arterial ECs in combination with *Tg(flt4:citrine)*^*hu7135*^, which is more strongly expressed in veins. In the distal fin, we observed continued vein-artery sprouting with vein ECs sprouting ventrally (Fig. 4A,A’’) before turning around and making a connection with the tail artery (Fig. 4B,blue arrowhead). This mode of sprouting continued as the fin expanded between 4 and 11 dpf (Supplementary Fig. 4). Thus, part of the fin vasculature forms through the repetitive addition of new artery-vein loops that emanate from a vein and connect to a pre-existing artery.

**Figure 4.**
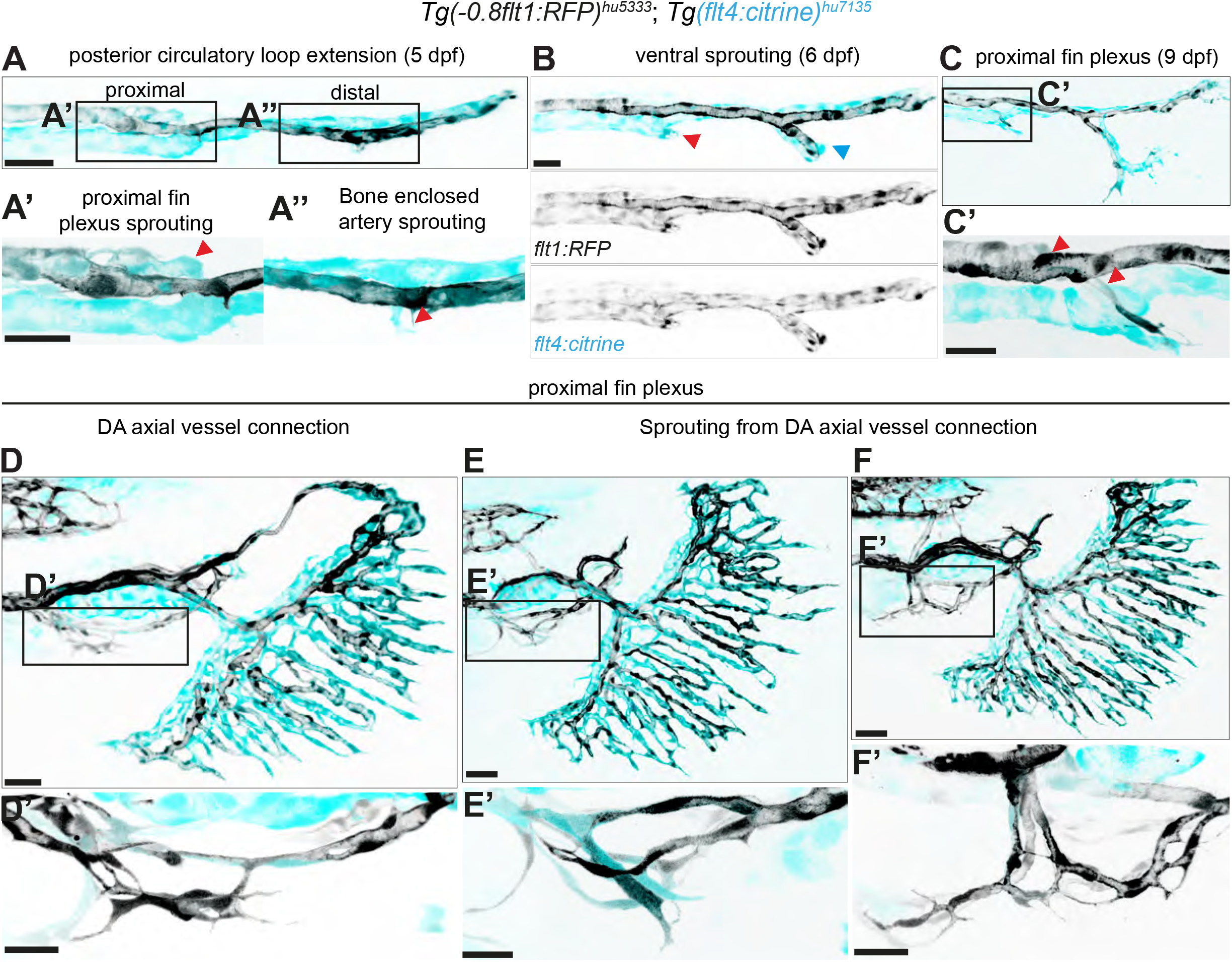
Proximal fin plexus and caudal fin arteries have distinct origins. Maximum intensity projections of confocal z-stacks of *Tg(−0*.*8flt1:RFP)*^*hu5333*^; *Tg(flt4: citrine)*^*hu7135*^ double transgenic fish labelling all arterial ECs (black) and venous cells (cyan) in lateral views, anterior to the left. **(A)** Axial circulatory loop at 5 dpf. Scale bar: 10 μm. Enlarged boxed regions show ventral sprouting from DA/PCV junction **(A’)** and axial vessels **(A’’)**. Scale bar: 4 μm. **(B, C)** Extension of posterior sprout and looping of axial sprouts at 6 dpf. Scale Bar: 15 μm. Boxed region in **(C’)** shows arterial sprout connection with the DA. Scale Bar: 5 μm. **(D-F)** Stages in the formation of proximal fin plexus. Scale Bar: 20 μm for **(D)**, 30 μm for **(E, F)**. Boxed regions in **(D’-F’)** shows the formation of the posterior plexus from the DA axial vessel connection. Scale Bar: 7 μm for **(D’, F’)**, 5 μm for **(E)**.

Of interest, we detected another, more proximally located, fin region that showed vascular expansion (Fig. 4A’-F’, red arrowheads). In this area, we did not observe the regular formation of well-patterned artery-vein loops, but rather the formation of a plexus. ECs within this plexus expressed varying levels of both the *flt1:RFP* and *flt4:citrine* transgenes when compared to the expanding arteries and veins in distal fin regions. Therefore, we can distinguish at least two distinct mechanisms that control fin blood vessel formation. In distal fin regions a more stereotyped process generates artery-vein loops that are quickly lumenized and carry flow, while in the proximal fin a less perfused vascular plexus forms. This plexus remodelled into 2 separate sets of arteries that ran on either side of the fish body at 3 wpf (Figure 5A-A’’). At this time point, another vascular plexus started to form at the base of the fanning fin vasculature, where a major vein and artery run in parallel in a dorsal to ventral fashion (Fig. 5B,B’). We named this vascular structure “interlaced plexus” to accommodate for its highly ramified morphology. Within this plexus, we detected new vascular connections that ran next to the pre-existing artery-vein loops and extended from proximal into distal fin regions (Fig. 5C, C’, pseudo-colored in red). They frequently displayed blind-ending tips that reached into more distal fin regions (Fig. 5C’, red arrowheads). Their expansion continued at 4 wpf, while being absent from the most distal regions of the fin (Fig. 5D,D’’). We found connections between these vessels and the bone enclosed arteries (Fig. 5D’’,yellow arrowheads). To characterize these vessels further, we used *Tg(pdgfrb:gal4ff)* ^*ncv24tg*^; *Tg(UAS:GFP)*^*nkuasgfp1a*^; *Tg(−0*.*8flt1:RFP)*^*hu5333*^ triple transgenic animals to label vascular mural cells (Fig. 5E,E’). This analysis revealed that vessels of the interlaced plexus readily contained mural cells, while their diameters were larger than those of bone enclosed arteries (Fig. 5F-G). They further lacked expression of the *flt4:citrine* transgene and showed lower levels of the arterial marker *flt1:RFP* (Fig. 5H). Analysis of blood flow patterns revealed a higher number of circulating red blood cells within the vessels of the interlaced plexus, when compared to bone enclosed arteries, as visualized through lower fluorescence intensity of injected qdots (Figure 5I-J’’, yellow arrowheads indicate blood cells). Together, our analysis reveals that fin vascular morphogenesis generates distinct types of blood vessels. An initial wave generates artery-vein loops that are positioned in relation to other morphological features, such as bone rays, followed by a second wave of angiogenic sprouting that generates less defined vascular plexuses. We can distinguish plexus vessels in terms of transgenic marker gene expression, diameters, and blood flow profiles. These successive processes furthermore occur in two different anatomical locations.

**Figure 5.**
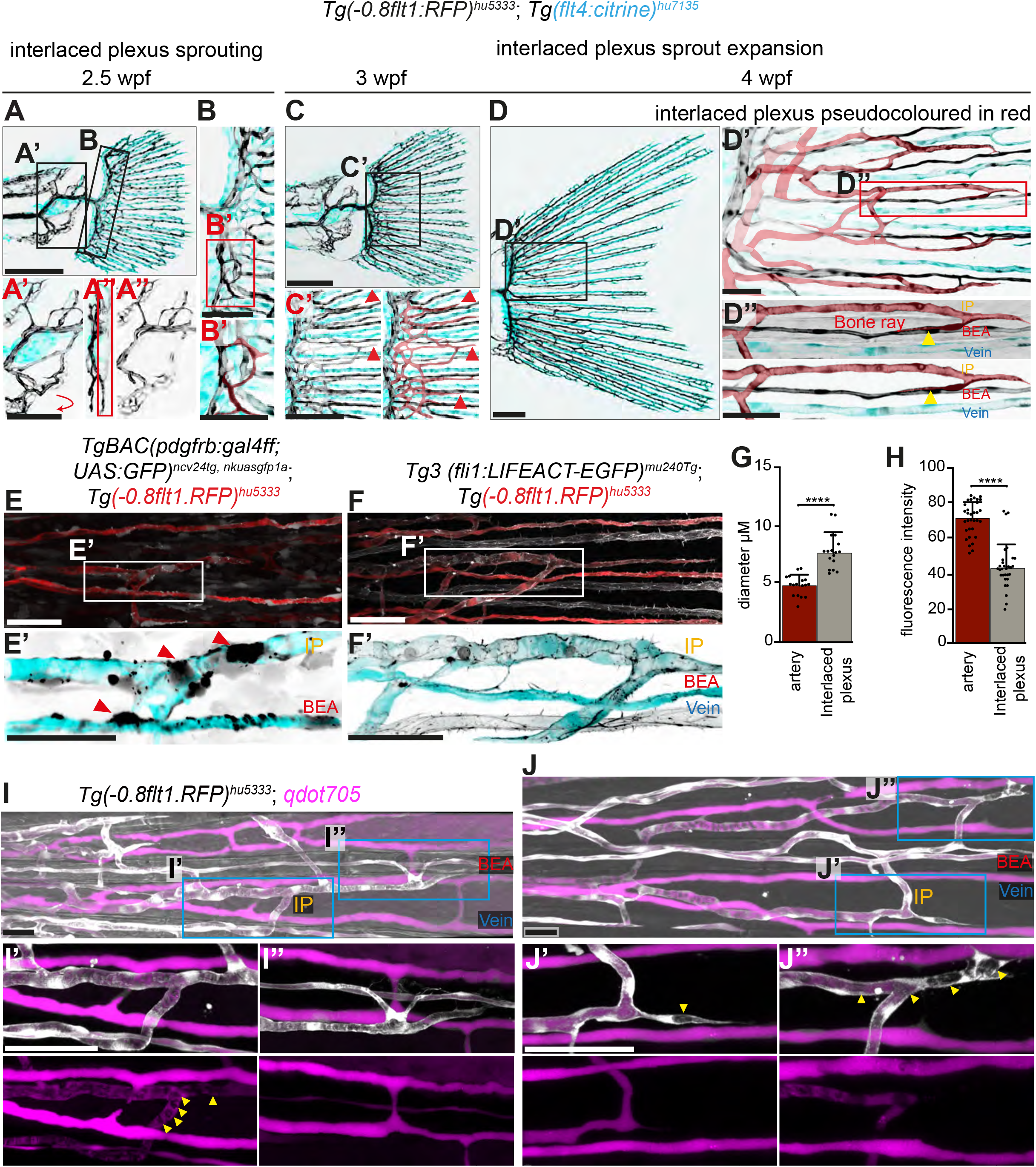
Formation of the interlaced plexus at the fin base. **(A-D)** Maximum intensity projections of confocal z-stacks of *Tg(−0*.*8flt1:RFP)*^*hu5333*^; *Tg(flt4: citrine)*^*hu7135*^ double transgenic fish labelling all arterial ECs (black) and venous cells (cyan) in lateral views anterior to the left. Sprouting and expansion of interlaced plexus from 2.5 wpf to 4 wpf. Scale Bar: 30 μm for **(A)**, 50 μm for **(C)**, 100 μm for **(E)**. Boxed regions enlarged in **(A’** and **A’’)** show the bifurcated proximal fin plexus. Scale Bar: 20 μm. Boxed region enlarged in **(B, B’)** shows the formation of initial loops of the interlaced plexus at the fin base at 2.5 wpf (pseudo-coloured in red in **B’)**. Scale bar: 10 μm for **(B)**, 5 μm for **(B’)**. Boxed region enlarged in **(C’-D’’)** shows expansion of interlaced plexus (pseudo-coloured in red.) Note blunt ends **(C’**, red arrowheads) and connections with bone enclosed arteries **(D’’**, yellow arrowheads). Scale bar: 10 μm for **(C’)**, 20 μm for **(D’)** and 15 μm for **(D’’). (E)** Maximum intensity projections of confocal z-stacks of *Tg(−0*.*8flt1:RFP)*^*hu5333*^; *TgBAC(pdgfrb:gal4ff;UAS:GFP)*^*ncv24tg, nkuasgfp1a*^ double transgenic fish labelling all arterial ECs (red/cyan) and mural cells (white/black) in lateral views, anterior to the left. Boxed region enlarged in **(E’)** shows mural cells on interlaced plexus and bone enclosed arteries (red arrowheads) Scale Bar: 10 μm for **(E)** and 5 μm for **(E’). (F)** Maximum intensity projections of confocal z-stacks of *Tg(−0*.*8flt1:RFP)*^*hu5333*^; *Tg3(fli:LIFEACT-EGFP)*^*mu240Tg*^ double transgenic fish labelling all arterial ECs (red/cyan) and all ECs (white/black) in lateral views anterior to the left at 4 wpf. Scale Bar: 20 μm for **(F)** and 7 μm for **(F’). (G)** Diameter of bone-enclosed artery and interlaced plexus. **(H)** Fluorescence intensity of bone-enclosed artery and interlaced plexus. **(I-J’’)** Maximum intensity projections of confocal z-stacks of *Tg(−0*.*8flt1:RFP)*^*hu5333*^; *qdot705* injections labelling arterial ECs (white) and blood vessel lumina (magenta). Yellow arrowheads indicate red blood cells in the interlaced plexus. Scale Bar: 20 μm for **(I-J)**, 7 μm for **(I’, I’’)**, 5 μm for **(J’, J’’)**. Paired t test. ***P=0.0001, ****P=<0.0001. n=17 segments from three individual fish for diameter measurements of IP and BEA, data points indicate individual segments. n=32 cells for BEA and n=30 cells for IP from three individual fish for fluorescent intensity measurements, data points indicate intensity values of individual cells.

### Sprouting from veins generates new veins to form vein-artery-vein triads at later developmental stages

At around 4 wpf, we could divide the tail fin vasculature into a dorsal and ventral lobe, with their plane of symmetry running through the centre of the fin (Fig. 6A-B’’). Thus, in the dorsal lobe, the arteries of individual fin rays came to lie dorsally to their respective veins, while in the ventral lobe, the arteries ran ventrally. At this stage, we observed a third morphogenetic process that we termed “secondary vein-sprouting”. This process led to the transformation of the initial artery-vein loops into triads of medially located arteries flanked by two veins (Fig. 6C). Secondary vein sprouting initiated at the centre of the fin ray, distally of the nascent interlaced plexus. Here, sprouts from the pre-existing first vein extended to form secondary veins (Fig. 6C, blue arrowhead). Connections from the first vein either crossed the bone enclosed artery, giving rise to secondary veins, or connected to the artery (Supplementary Fig. 5). Thus, in contrast to the initial angiogenic sprouting, where veins gave rise to arteries, secondary vein sprouting exclusively generates veins from pre-existing veins.

**Figure 6.**
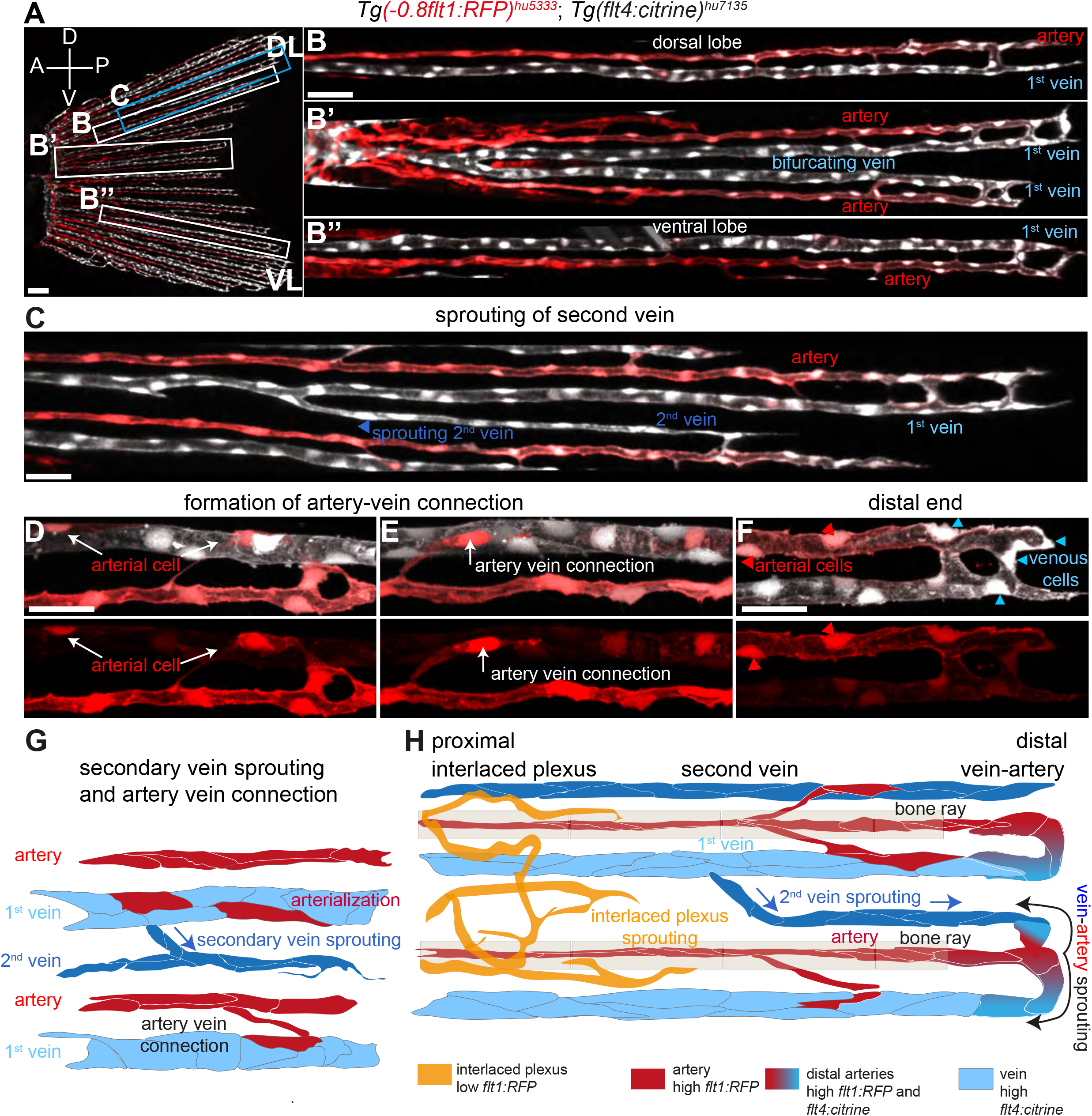
The fin vascular network expands through vein derived sprouting of new veins and arteries. **(A)** Maximum intensity projections of confocal z-stacks of *Tg(−0*.*8flt1:RFP)*^*hu5333*^; *Tg(flt4: citrine)*^*hu7135*^ double transgenic fish at 3 wpf labelling all arterial ECs (red) and venous cells (white) in lateral views anterior to the left. Scale bar: 20 μm. **(B)** Dorsal lobe, first vein forms below the bone enclosed artery. **(B’)** Mid lobe, vein bifurcates forming the first vein above the bone enclosed artery. **(B’’)** Ventral lobe, first vein forms above bone enclosed artery. Scale bar: 30 μm. **(C)** Growth of second vein through lateral sprouting from first vein. Scale bar: 30 μm. **(D, E)** Formation of artery – vein connections. Activation of arterial marker in single venous ECs (white arrows). Scale bar: 10 μm. **(F)** Distal end of caudal fin blood vessels. Note the presence of venous ECs (blue arrowheads) at the tip of newly forming arteries (red arrowhead). Scale bar: 10 μm. **(G)** Schematic representation of second vein sprouting and artery – vein connections. **(H)** Schematic representation of caudal vein vasculature at 3 wpf depicting four distinct pattens of new blood vessel formation occurring simultaneously.

We also observed the establishment of new connections between arteries and veins along the length of the fin vasculature. Here, ECs within veins started to express the *flt1:RFP* transgene (Fig. 6D) before connecting to arteries (Fig. 6E). Of note, at the most distal end of the fin, vein cells continued to generate arteries through “vein-artery sprouting” (Fig. 6F). Thus, as the fin vasculature expands, vein cells at the distal tip simultaneously generate new vein and arterial cells, while in more proximal regions, these processes appear to be uncoupled. Here, veins generate new vein cells through “secondary vein sprouting” without generating arterial cells. In different locations, cells in the first vein differentiate into arterial cells without making new vein cells to establish new artery-vein connections. This network of blood vessels is being complemented by a plexus of interlaced arteries in more proximal locations that forms through distinct mechanisms. Together, at this developmental time point, four simultaneously occurring, distinct morphogenetic processes expand the tail fin vasculature along the proximal-distal axis (interlaced plexus sprouting, vein arterialization, second vein sprouting and vein-artery sprouting, summarized in Fig. 6G,H).

### Intussusceptive angiogenesis expands the arterial network

After the establishment of the initial fin vasculature and artery-vein connections, we observed another vein-derived expansion of the arterial network through intussusceptive angiogenesis (Fig. 7A-D). ECs at the outer edges of distally located veins started to downregulate the *flt4:citrine* transgene (Fig. 7C,blue arrowheads), while increasing *flt1:RFP* expression (Fig. 7C,red arrowheads). Of note, we detected cytoplasmic pillars, characteristic of intussusceptive angiogenesis (De Spiegelaere et al., 2012) (Fig. 7C’,demarcated by red lines). Ultimately, *flt1:RFP* expressing ECs detached from their vein of origin and coalesced into small diameter arteries that now ran adjacent to the pre-existing vein and connected to other artery-vein connections (Fig. 7D,red arrowheads, summary in Fig. 7G). Thus, intussusceptive angiogenesis is the fifth morphogenetic process we detected in the expanding fin vasculature.

**Figure 7.**
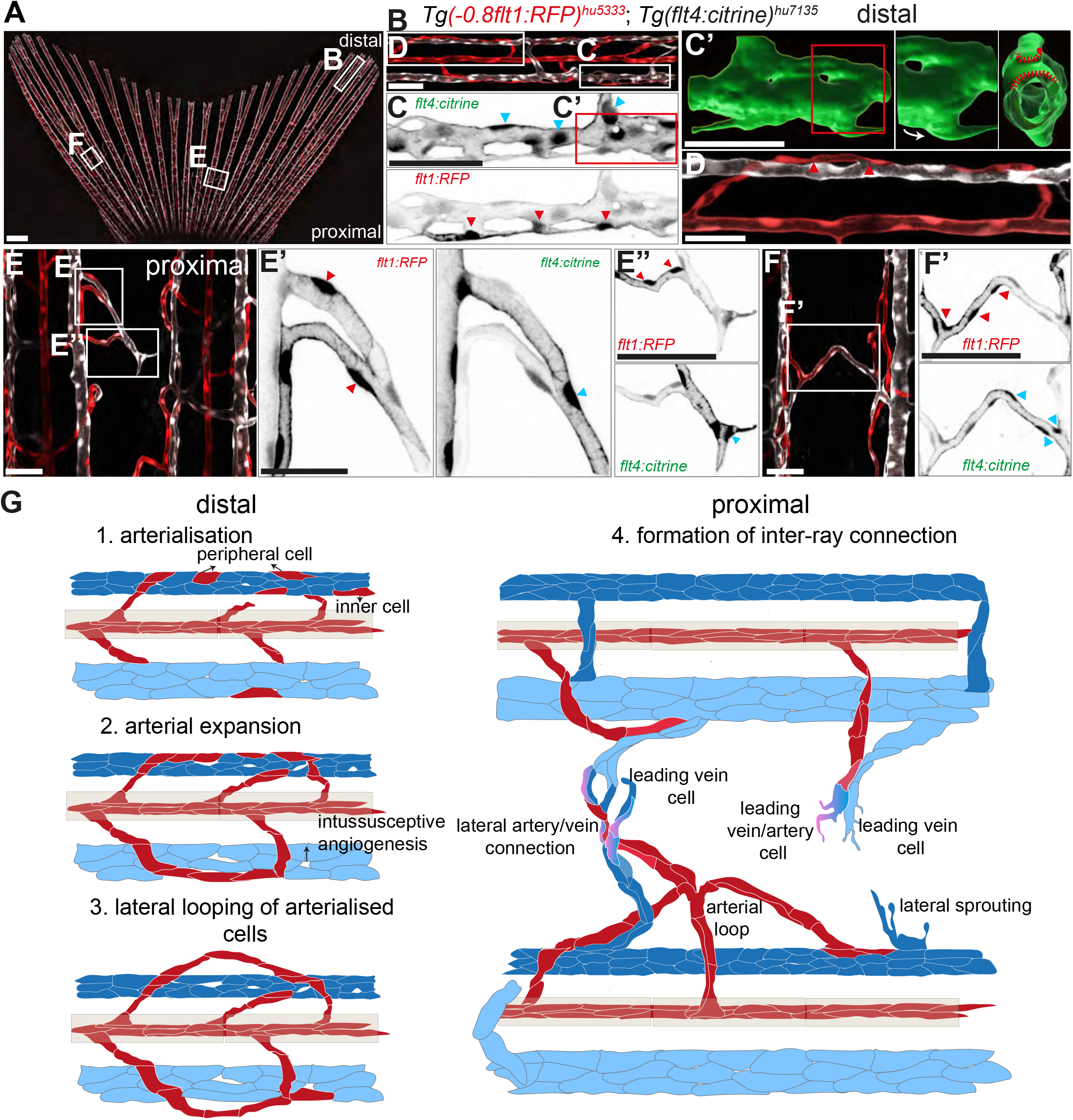
Inter-ray connections are formed by lateral sprouting and intussusceptive angiogenesis. **(A)** Maximum intensity projections of confocal z-stacks of *Tg(−0*.*8flt1:RFP)*^*hu5333*^; *Tg(flt4: citrine)*^*hu7135*^ double transgenic fish labelling all arterial ECs (red) and venous cells (white). Caudal fin of 1.5-month-old fish. Scale Bar: 300 μm. **(B)** Distal end of caudal fin vasculature. Scale bar: 20 μm. **(C)** Arterialization in ECs of veins, blue arrowheads indicate venous ECs, red arrowheads indicate arterialized ECs. Scale bar: 5 μm. **(C’)** 3D reconstruction of boxed region in **(C)** shows the formation of pillars in veins (red dashed lines). **(D)** Connection between bone enclosed artery and peripheral arterial cells (red arrowheads). Scale Bar: 10 μm. **(E)** Lateral connections forming in the proximal regions of the fin. Scale Bar: 20 μm. Boxed regions enlarged in (**E’** and **E’’**) show lateral sprouts originating from veins. Note the presence of venous ECs occupying the leading edge of new sprouts (blue arrowheads), while arterial cells trail behind (red arrowheads). Scale Bar: 4 μm for **E’** and 5 μm for **E’’. (F, F’)** Fully formed inter ray connection between artery (red arrowheads) and vein (blue arrowheads). Scale Bar: 20 μm for **(F)** and 5 μm for **(F’). (G)** Stages in the formation of inter-ray connections. Distal: (1) arterialisation in inner and peripheral ECs of the vein, (2) expansion arterialised ECs and establishment of connection with bone enclosed arteries, (3) lateral looping of arterialised ECs. Proximal: (4) lateral sprouting from veins establishes connections with arterial loops and further extends to form inter-ray connections.

### Sprouting angiogenesis generates capillary-type connections between arteries and veins in proximal fin regions

In more proximal areas, we observed the formation of a more intricate network of smaller diameter interconnections between arteries and veins. Here, ECs at the tips of crossing sprouts more strongly expressed the *flt4:citrine* transgene when compared to the *flt1:RFP* transgene (Fig. 7E-E’’). Cells at the base of the sprouts either connected to a larger vein and expressed *flt4:citrine* or to a smaller artery and expressed *flt1:RFP* (Fig. 7E-E’’). In established connections and when moving from the arterial towards the venous side of the vascular tree, we observed a gradual transition from *flt1:RFP* transgene expression (Fig. 7F,F’,red arrowheads) to *flt4:citrine* transgene expression (Fig. 7F,F’,blue arrowheads). Thus, the later stages of fin vascular development are characterized by an elaboration of the initially laid down framework of alternating arteries and veins (Fig. 7G). Distally located vessels mainly establish arterial side branches that connect to veins within a given fin ray through intussusceptive angiogenesis, while more proximal fin regions generate additional connections through sprouting angiogenesis that span between fin rays (Summary of all morphogenetic processes in Figure 8).

**Figure 8.**
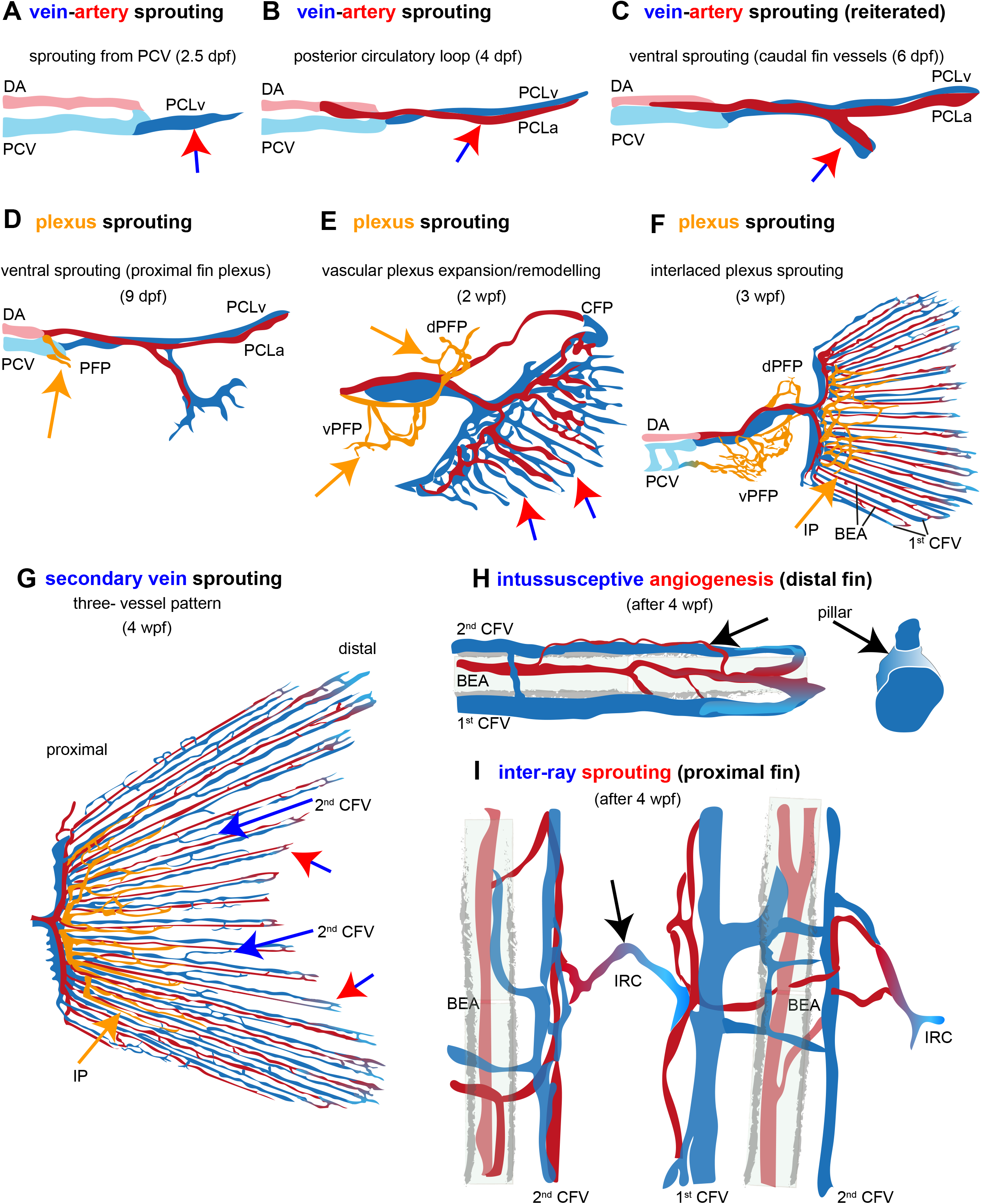
Schematic representation of caudal fin vascular development. **(A)** Initiation of caudal fin development at 2.5 dpf from the PCV in a process we term vein-artery sprouting, indicated by red-blue arrow. **(B)** Posterior circulatory loop consists of PCLa connected to DA and PCLv connected to PCV. **(C)** Ventral sprouting from PCLv. PCLv sprouts and loops back to PCLa to form the ventral loop. **(D)** Ventral sprouting from DA/PCV junction initiates at 9 dpf to form the PFP in a process we term plexus sprouting. **(E)** at 2 wpf PFP expands to form dPFP and vPFP, CFP expands into growing fin. **(F)** At 3 wpf the CFP remodels into a two-vessel pattern with BEA and 1^st^ CFV. An IP forms. **(G)** Secondary vein sprouting establishes a three-vessel pattern. IP expands. **(H)** Intussusceptive angiogenesis in distal fin regions generates new arterial side branches from veins (black arrow). **(I)** Inter-ray connections are formed in the proximal regions of the fin (black arrow). Abbreviations: DA: dorsal aorta, PCV: posterior cardinal vein, PCLv: posterior circulatory loop vein, PCLa: posterior circulatory loop artery, PFP: proximal fin plexus, vPFP: ventral proximal fin plexus, dPFP: dorsal proximal fin plexus, CFP: caudal fin plexus, IP: interlaced plexus, BEA: bone enclosed artery, 1^st^ CFV: first caudal fin vein, 2^nd^ CFV: second caudal fin vein, IRC: inter ray connection.

## Discussion

During embryonic development, every organ of the body needs to be vascularized. This entails the tight coupling of tissue development with vascular growth (Augustin and Koh, 2017). Previous studies have shown that specialized vascular beds exist within distinct organs and that blood vessels can also direct organ development (Gomez-Salinero *et al*., 2021; Lammert et al., 2001; Rafii *et al*., 2016; Stenman et al., 2008). Blood vessels furthermore align with anatomical structures, such as bones and nerves (Bump *et al*., 2022; Larrivee et al., 2009). Previous studies identified hard-wired genetic mechanisms that lead to the formation of a stereotyped vasculature, for example within the trunk of the zebrafish embryo (Childs et al., 2002; Hogan and Schulte-Merker, 2017; Isogai et al., 2001; Siekmann and Lawson, 2007), the hindbrain (Bussmann et al., 2011; Fujita et al., 2011), or the pectoral fins (Paulissen *et al*., 2022). However, we previously showed that the hindbrain vasculature in individual wild-type embryos differed in terms of blood vessel connectivity that generated a considerable degree of variability, likely due to the regulation of *cxcr4a* expression by blood flow (Bussmann *et al*., 2011). The same is true for the trunk vasculature after the later occurring sprouting of secondary veins that establish a re-wired arterial-venous network (Geudens et al., 2019; Weijts et al., 2018). Thus, while some aspects of blood vessel sprouting appear to follow precise guidance cues, other processes are more flexible. This notion of variability is further underscored by the discovery of blood vessel pruning, where unnecessary initial connections are being removed, leading to an optimization of blood flow patterns (Chen et al., 2012; Franco et al., 2015; Kochhan et al., 2013; Korn and Augustin, 2015; Lenard et al., 2015). While many of the genetic players orchestrating vascular growth and blood vessel sprouting, such as VEGF, Notch, WNT and BMP signalling have been identified (Adams and Alitalo, 2007; Eelen *et al*., 2020; Potente et al., 2011; Red-Horse and Siekmann, 2019; Schuermann et al., 2014), our insights into the mechanisms controlling the stepwise assembly of the vasculature of a given organ and how these might regulate vascular variability have remained limited.

To investigate the different morphogenetic processes that are required to shape the adult vasculature of an organ and to understand how these processes might build upon each other, we decided to study the developing tail fin vasculature of zebrafish embryos over a time frame of 4 weeks. We chose this vascular bed, as the fin is relatively two-dimensional and using transparent zebrafish allows for the imaging of vascular network assembly at later developmental time points (Xu *et al*., 2014). Our analysis revealed that the initial vascular loop in the fin formed through angiogenic sprouting from the posterior cardinal vein (PCV). In this process, vein-derived ECs relied on VEGF to turn on arterial marker gene expression and change their direction of migration, ultimately turning towards the dorsal aorta in a Notch and *cxcr4a*-dependent manner. This sequence of events that we now termed vein-artery sprouting was reiterated during the subsequent outgrowth of the fin vasculature into the expanding fin fold, generating an artery-vein loop within each fin ray. We previously identified the same genetic program during eye blood vessel development in zebrafish embryos and in the mouse retina, describing a process through which vein-derived angiogenic ECs form new arteries (Hasan *et al*., 2017; Pitulescu et al., 2017; Xu *et al*., 2014). Similar vein to artery conversions have been described for the coronary arteries (Red-Horse et al., 2010; Su et al., 2018), in early mouse embryos (Hou et al., 2022) and further in the mouse retina (Lee et al., 2021; Park et al., 2021). In the pectoral fin, two vein sprouts initially emerge from the common cardinal vein and fuse along the rim of the endoskeletal disc (Paulissen *et al*., 2022). Later, a new sprout grows from this vein loop towards the dorsal aorta. Thus, also in the pectoral fin, veins are the source for new ECs, followed by artery specification and migration of a new sprout towards a pre-existing artery. This sprouting depended on Notch signalling (Paulissen *et al*., 2022) and we speculate that sprouting towards the dorsal aorta will also depend on *cxcr4a* signalling. Together, these findings underscore the conserved nature of this mode of blood vessel formation that generates perfused artery-vein loops, in which arteries and veins can be distinguished by virtue of marker gene expression, diameter, and blood flow (Red-Horse and Siekmann, 2019). However, this configuration of direct artery-vein loops allows arterial flow patterns to be directly transmitted into venous blood vessels, as we do not observe capillary beds at this time point in the fin. Therefore, the initial expansion of the fin vasculature generates vascular patterns consisting of what is traditionally viewed as arterio-venous shunts (Pries et al., 2010).

Curiously, we then observed a shift in the mode of blood vessel sprouting. New sprouts again emerged from pre-existing veins, but exclusively generated new veins instead of arteries in a process we call secondary vein sprouting. These veins migrated in proximal and distal directions along the bone ray and either connected with a pre-existing vein or the bone-enclosed artery in the most distal fin region, effectively transforming the direct artery-vein connections into a pattern, where one artery now drains into two veins. Therefore, this mode of angiogenic sprouting resolved the initially present arterio-venous shunts. A similar vessel pattern is also observed in pectoral fins (Paulissen *et al*., 2022). This switch in angiogenic sprouting behaviour thus appears to be a fundamental process to ensure appropriate flow patterns through a later expansion of the venous pole of the vasculature. It will be important in the future to understand how this switch in angiogenic sprouting is achieved and whether it also occurs in other vascular beds. Previous studies implicated bone morphogenetic proteins (BMPs) in controlling vein sprouting in the trunk of zebrafish embryos (Wiley et al., 2011). In this setting, a first wave of angiogenic sprouting generates intersegmental blood vessels that are connected to the dorsal aorta in a process dependent on VEGF and Notch signalling (Hogan and Schulte-Merker, 2017; Nasevicius et al., 2000; Siekmann and Lawson, 2007), while in this setting *cxcr4a* signalling is dispensable (Siekmann et al., 2009). Subsequently, secondary sprouts re-wire this initial artery-only network into arteries and veins (Geudens *et al*., 2019; Weijts *et al*., 2018). It is therefore likely that BMPs might also control vein-generating sprouting in the tail fin, as BMPs have similarly been implicated in vein formation in other contexts (Neal et al., 2019). BMP signalling is strongly influenced by blood flow (Baeyens et al., 2016a; Baeyens et al., 2016b; Vion et al., 2018; Zhou et al., 2012). It will be of interest to investigate whether changing flow patterns in the tail fin might alter signalling within ECs to orchestrate the switch from angiogenic sprouting that generates arteries from veins to a program that exclusively generates veins. Together, our results demonstrate that two distinct angiogenic programs exist that allow for the expansion of the venous pole of the vasculature after the initial establishment of artery-vein loops.

We observed an additional set of blood vessels that formed at the base of the fin through a process we term plexus sprouting. These could be distinguished in terms of their diameters, transgene expression and blood flow patterns. We named these blood vessels the interlaced plexus. It is currently unclear from which blood vessels these ECs emerge, as they do not show expression of the vein-enriched *flt4:citrine* transgene. They further express lower levels of the arterial-enriched *flt:RFP1* transgene. Of interest, low levels of *flt1:RFP* expression were recently reported in lymphatic ECs that give rise to blood vessels in the anal fin (Das *et al*., 2022). It might therefore be conceivable that the cells of the interlaced plexus have a similar lymphatic origin, albeit the absence of *flt4:citrine* expression argues against this possibility. ECs forming veins in the pectoral and tail fin vasculature also express the lymphatic marker *mrc1a:egfp* (Paulissen et al., 2022). How these gene expression patterns reflect functional properties of the investigated blood vessels, and their origin remains to be determined.

Lastly, we observed two distinct modes of blood vessel formation that expanded arterial collateral vessels in proximal and distal fin regions, respectively. In more distal regions, we observed intussusceptive angiogenesis in veins (De Spiegelaere *et al*., 2012). This process occurs through the insertion of tissue pillars within blood vessels that subsequently expand, leading to a splitting of a blood vessel into two individual vessels (Burri and Djonov, 2002). Intussusceptive angiogenesis frequently occurs after sprouting angiogenesis and has been observed in the caudal vein plexus of zebrafish embryos (Karthik et al., 2018). Despite the detailed description of the processes involved in intussusceptive angiogenesis, changes in gene expression patterns within ECs undergoing intussusceptive angiogenesis have not been detected on the cellular level. We now observed that individual ECs within veins started to express the arterial marker *flt1:RFP* during intussusceptive angiogenesis and coalesced into separate blood vessels made up of *flt1:RFP* expressing ECs that connected to arterial side branches. These branches did not carry red blood cells, and we can only speculate as to their function. They might play a role during blood vessel regeneration to serve as collaterals that would restore blood flow after the main artery has been severed. We previously observed that these interconnections were enlarged during artery regeneration in animals lacking mural cells (Leonard et al., 2022), with a role of vascular mural cells being described during intussusceptive angiogenesis (Burri et al., 2004). Further studies will be required to fully elucidate their role and what mechanisms control the observed changes in gene expression in ECs undergoing intussusceptive angiogenesis, with changes in hemodynamics playing a potential role (Styp-Rekowska et al., 2011).

In more proximal regions of the tail fin, we observed inter-ray sprouting angiogenesis from veins. Sprouts regularly crossed the space between two adjacent fin rays. They expressed varying levels of arterial and venous marker genes that either formed sharp boundaries or displayed more gradual changes from the arterial towards the venous pole of the vasculature, as also recently described for capillaries forming during mouse embryogenesis (Hou *et al*., 2022). Therefore, in proximal fin regions small blood vessels resembling a capillary bed form at later stages of fin vascular expansion.

Together, our longitudinal analysis of the development of the zebrafish tail fin vasculature illustrates how blood vessels assemble in a stepwise manner to irrigate a mature organ. Our results furthermore show how different morphogenetic processes are either reiteratively used, as for the generation of arteries from veins, or build upon each other, as in the case for the expansion of the venous vasculature to transform arterio-venous shunts into a configuration where one artery drains into two veins. How these underlying principles organize the formation of blood vessels in other organs, as well as the signalling pathways controlling these steps, and the mechanisms through which they are being activated remains to be determined.

## Acknowledgements

We would like to thank Reinhild Bussmann, Mona Finch-Stephen, Bill Vought and Nadine Greer for excellent fish care. We would also like to thank Mike Pack for critical reading of the manuscript.

## Contributions

E.V.L. and S.S.H. performed experiments. A.F.S. conceptualized and supervised the work. E.V.L. and A.F.S. wrote the manuscript.

## Funding

This work was supported by grants from the Max-Planck-Society, the Deutsche Forschungsgemeinschaft (DFG SI 1374/5-1 and SI 1374/6-1) and from the National Heart, Lung and Blood Institute (R01HL152086) awarded to A.F.S.. E.L. was partly supported by funds from the Cluster of Excellence Cells in Motion (CiM) of the University of Münster.

## Materials and Methods

### Zebrafish husbandry and strains

Zebrafish embryos were maintained in 1X E3 under recommended husbandry conditions (Westerfield, 1993). Embryos were transferred to tanks in the zebrafish facility and raised until further analysis. Previously described zebrafish lines were *Tg(−0*.*8flt1:RFP)*^*hu5333*^ (Bussmann et al., 2010), *TgBAC(pdgfrb:gal4ff)*^*ncv24*^ (Ando et al., 2016), *Tg(TP1bglob:VenusPEST)*^*s940*^ (Ninov et al., 2012), *Tg(TP1bglob:H2B-mCherry)*^*s939*^ (Ninov *et al*., 2012), *Tg3(fli1:LIFEACT-EGFP)*^*mu240*^ (Hamm et al., 2016), *Tg(fli1a:nEGFP)*^*Y7*^ (Roman et al., 2002), *Tg(gata1:dsRed)*^*sd2*^ (Traver et al., 2003), *TgBAC(cxcr4a:YFP)*^*mu104*^ (Xu *et al*., 2014), *Tg (flt4:citrine)*^*hu7135*^ (van Impel et al., 2014), *Tg(kdrl:EGFP)*^*s843*^ (Jin et al., 2005), *kdrl*^*hu5088*^ (Bussmann *et al*., 2010), *vegfaa*^*mu128*^ (Lange et al., 2022). Some of the transgenic lines were kept in a *casper* (*roy, nacre*) double mutant background (White et al., 2008) to increase clarity for imaging due to loss of pigment cells. All animal experiments were performed in compliance with the relevant laws and institutional guidelines and were either approved by local animal ethics committees of the Landesamt für Natur, Umwelt und Verbraucherschutz Nordrhein-Westfalen or in accordance with protocols approved by the University of Pennsylvania Institutional Animal Care and Use (number 806819). Zebrafish veterinary care was performed under the supervision of the University Laboratory Animal Resources at the University of Pennsylvania.

### Time lapse imaging and angiography

Embryos (2-3 dpf) were embedded in 1% low melting point agarose containing 168mg/l tricaine in a glass bottom dish. Embryos were imaged for the desired period of time with 20 min imaging intervals. A heated microscope stage set to 28.5°C was used to maintain the appropriate temperature. Larvae from 4 dpf to 4 wpf were anaesthetized in tricaine and later embedded in 1% low melting point agarose. For imaging of 1.5-month-old fish, caudal fins were amputated from anaesthetized fish and embedded in 1% low melting point agarose for imaging. Angiography was performed by injecting qdots 705 into the hearts of 4 wpf fish.

### Inhibitor treatment

SU5416 (Millipore Sigma) was dissolved in DMSO and stored at -20°C. Embryos were dechorionated and treated with 2.5 µM of SU5416 or 0.1 % DMSO in E3 media beginning at 48 hpf until 72 hpf. The drug solution was refreshed every 12 hrs. Embryos were incubated in E3 media containing 0.003% phenylthiourea to inhibit pigmentation.

### Confocal microscopy

Fluorescent confocal images were acquired using a Zeiss LSM 780 (objective lens: 20x Plan Apo NA 0.80) or Leica SP8 (objective lens: HC PL Fluotar 20x/0.50 or HCX APO L 63x). All images shown are representative of the results obtained for each group and experiment.

### Image processing

Zeiss ZEN software or Leica LAS-X were used to stitch the acquired images. Imaris software (Oxford Instruments) was used to generate maximum intensity projections. 3D projections were made by creating surfaces in Imaris. ImageJ (NIH) was used to generate monochrome images of different fluorophores. Adobe Illustrator and Adobe Photoshop software were used to compile images and create schematics.

### Quantification and analysis

Nuclei numbers were manually counted using the Imaris software suite (Oxford Instruments) in blood vessels of approximately the same length. Diameters of interlaced plexus vessels and bone enclosed arteries was calculated by measuring at least 10 regions in segments of equal length. Fluorescence intensity was calculated using maximum intensity projections. Individual cells were marked, and the intensity values were obtained from ImageJ (NIH). Data were analysed using Prism 9 (Graphpad) and graphs were plotted with mean standard deviation (SD). p values <0.05 were considered statistically significant.

## Notes

### Competing Interest Statement

The authors have declared no competing interest.

