## Supplementary for "Temporally and regionally distinct morphogenetic processes govern zebrafish tail fin blood vessel network expansion"

**Supplementary Figure 1. Still images of additional movies showing vein-artery sprouting in wild-type and *cxcr4a$^{um}$* mutants.**

**(A,B)** Maximum intensity projections of confocal z-stacks of *Tg(fli1a:nEGFP)$^{y7}$*; *Tg (-0.8flt1:RFP)$^{hu5333}$* double transgenic embryos, where all EC nuclei are labelled in green, and all arterial ECs are labelled in red in wild-type embryos with time points indicated. Scale bar: 50 μm. **(C,D)** Maximum intensity projections of confocal z-stacks of *Tg(fli1a:nEGFP)$^{y7}$*; *Tg (-0.8flt1:RFP)$^{hu5333}$* double transgenic embryos, where all EC nuclei are labelled in green, and all arterial ECs are labelled in red in *cxcr4a$^{um20}$* embryos with time points indicated. Scale bar: 50 μm.

**Supplementary Figure 2. Formation of the first circulatory loop of fin vasculature.**

**(A, B)** Maximum intensity projections of confocal z-stacks of *Tg(fli1a:lifeactGFP)$^{mu240}$*; *Tg(-0.8flt1:RFP)$^{hu5333}$* double transgenic fish labelling all ECs (white) and arterial cells (red). Sprouting and extension from the PCV. Scale bar: 10 μm. **(C)** Arterialization in cells located ventral to the vein sprout (red arrowheads). Scale bar: 10 μm. **(D)** Reverse migration of arterialized ECs (red arrowheads). Scale bar: 10 μm. **(E)** Anastomosis of arterialized ECs with the DA (red arrowheads). Scale bar: 10 μm. **(E')** Anastomosis of newly formed artery to the ventral surface of the DA (red arrowheads). Scale bar: 5 μm. **(F)** Maximum intensity projections of confocal z-stacks of *Tg(fli1a:lifeactGFP)$^{mu240}$*; *Tg(gata1:dsRed)$^{sd2}$* double transgenic fish labelling all ECs (white) and red blood cells (red). Sprouting from the PCV (red arrowhead). Scale bar: 10 μm. **(G)** Lumenization and oscillatory blood flow (double red arrow), yellow arrowheads show red blood cells. Scale bar: 10 μm. **(H)** Sprout extension (yellow arrowheads indicate red blood cells). Scale bar: 10 μm. **(I)** Reverse migration of ECs and anastomosis with the DA. Note change in blood flow pattern (red arrows in bottom image). Scale bar: 10 μm. Boxed region enlarged in **(I')** shows lumenized vessels with blood cells (yellow arrowheads) and blood flow (red arrows in bottom image). Scale bar: 5 μm.

**Supplementary Figure 3. Still images from time-lapse movie showing *cxcr4a* expression in ECs with Notch pathway activation and failure of vein-artery sprouting in *vegfaa*$^{mu128}$ mutants.**

**(A)** Maximum intensity projections of confocal z-stacks of *Tg(fli1a:nEGFP)*$^{y7}$; *TgBAC(cxcr4a:YFP)*$^{mu104}$; *Tg(TP1:H2B-mcherry)*$^{s939}$ triple transgenic fish labelling all EC nuclei (blue), *cxcr4a* expressing cells (green) and cells with Notch pathway activation (red) in lateral views anterior to the left. White arrowheads indicate arterial ECs with activated Notch signalling. Scale bar: 20 µm. Boxed regions are enlarged. Scale bar: 4 µm. **(B)** Maximum intensity projections of confocal z-stacks of *Tg(fli1a:nEGFP)*$^{y7}$; *Tg(kdrl:Hsa.HRAS-mCherry)*$^{s916}$; double transgenic wild-type control fish labelling all EC nuclei (green) and EC membranes (red) in lateral views anterior to the left. Scale bar: 15 µm. **(C)** Maximum intensity projections of confocal z-stacks of *Tg(fli1a:nEGFP)*$^{y7}$; *Tg(kdrl:Hsa.HRAS-mCherry)*$^{s916}$; double transgenic *vegfaa*$^{mu128}$ mutant fish labelling all EC nuclei (green) and EC membranes (red) in lateral views anterior to the left. Scale bar: 15 µm. Note absence of vein-artery sprouting in *vegfaa*$^{mu128}$ fish.

**Supplementary Figure 4. Reiterative vein-artery sprouting during ventral growth and expansion of the caudal fin vascular network.**

**(A)** Maximum intensity projections of confocal z-stacks of *Tg(fli1a:nEGFP)*$^{y7}$; *Tg(-0.8flt1:RFP)*$^{hu5333}$ double transgenic embryos, where all EC nuclei are labelled in green, and all arterial ECs are labelled in red. Formation of first ventral sprout between 4 dpf and 1 wpf. Arrowheads indicate growing ventral sprout. Scale bars: 50 µm. Schematics represent the ventral sprouting process. White arrows denote blood flow direction. Numbers represent the individual embryos (or fish) analysed for each developmental stage. **(B)** Expansion of the primary ventral sprout to form more vascular branches between 8 dpf and 11 dpf. Arrowheads indicate branching points. Schematics represent expansion of the vascular tree. White arrows denote blood flow direction. Scale bars: 50 µm. Numbers represent the individual embryos analysed for each developmental stage. **(C)** Still images at different time points indicating formation of second arterial connections (arrowheads) between 8 dpf and 11 dpf. Scale bar: 50 µm. Schematics represent expansion of the vascular tree. White arrows denote blood flow direction. Numbers represent the individual embryos analysed for each developmental stage.

**Supplementary Figure 5. Morphogenesis of caudal fin vessels at 3-4 wpf.**

**(A)** Maximum intensity projections of confocal z-stacks of *Tg(fli1a:H2B-mcherry)*$^{uq37bh}$; *Tg fli1a:lifeactGFP)*$^{mu240}$. Three different types of connections are formed: **(B)** vein – vein **(C)** artery – vein **(D)** vein – artery. Scale bar: 10 μm. Boxed regions are enlarged. Scale bar: 5 μm.

**Supplementary Movie 1. ECs sprouting from the PCV form the first posterior circulatory loop.**

Maximum intensity projections of confocal z-stacks of *Tg(fli1a:nEGFP)*$^{y7}$; *Tg(-0.8flt1:RFP)*$^{hu5333}$ double transgenic embryos, where all EC nuclei are labelled in green, and all arterial ECs are labelled in red. Time-lapse imaging reveals endothelial cells sprouting from PCV and reverse migration of ECs expressing *flt1:RFP* to form the posterior circulatory loop. Scale bar: 50 μm.

**Supplementary Movie 2. ECs sprouting from the PCV form the first posterior circulatory loop.**

Maximum intensity projections of confocal z-stacks of *Tg(fli1a:nEGFP)*$^{y7}$; *Tg(-0.8flt1:RFP)*$^{hu5333}$ double transgenic embryos, where all EC nuclei are labelled in green, and all arterial ECs are labelled in red. Time-lapse imaging reveals endothelial cells sprouting from PCV and reverse migration of ECs expressing *flt1:RFP* to form the posterior circulatory loop. Scale bar: 50 μm.

**Supplementary Movie 3. ECs sprouting from the PCV form the first posterior circulatory loop.**

Maximum intensity projections of confocal z-stacks of *Tg(fli1a:nEGFP)*$^{y7}$; *Tg(-0.8flt1:RFP)*$^{hu5333}$ double transgenic embryos, where all EC nuclei are labelled in green, and all arterial ECs are labelled in red. Time-lapse imaging reveals endothelial cells sprouting from PCV and reverse migration of ECs expressing *flt1:RFP* to form the posterior circulatory loop. Scale bar: 50 µm.

**Supplementary Movie 4. Non lumenized caudal fin sprout at 2.5 dpf.**

Time-lapse confocal imaging of *Tg(kdrl:EGFP)*$^{s843}$; *Tg(gata:dsRED)*$^{sd2}$ double transgenic embryos, where all ECs are labelled in green, and all erythrocytes are labelled in. Note the non lumenized sprout at 2.5 dpf. Scale Bar: 20 µm.

**Supplementary Movie 5. Lumenized caudal fin sprout displays oscillatory blood flow.**

Time-lapse confocal imaging of *Tg(kdrl:EGFP)*$^{s843}$; *Tg(gata:dsRED)*$^{sd2}$ double transgenic embryos, where all ECs are labelled in green, and all erythrocytes are labelled in red in lateral views anterior to the left, 56hpf. Note oscillatory blood flow. Scale Bar: 20 µm.

**Supplementary Movie 6. Artery-directed reverse migration of ECs is defective in *cxcr4a*$^{um20}$ mutants.**

Maximum intensity projections of confocal z-stacks of *Tg(fli1a:nEGFP)*$^{y7}$; *Tg(-0.8flt1:RFP)*$^{hu5333}$ double transgenic *cxcr4a*$^{um20}$ mutants, labelling all EC nuclei (green) and all arterial ECs (red). White arrowheads with numbers mark individual ECs. Note accumulation of arterialised ECs at the tip of the sprout and absence of reverse migration.

**Supplementary Movie 7. Artery-directed reverse migration of ECs is defective in *cxcr4a*$^{um20}$ mutants.**

Maximum intensity projections of confocal z-stacks of *Tg(fli1a:nEGFP)*$^{y7}$; *Tg(-0.8flt1:RFP)*$^{hu5333}$ double transgenic *cxcr4a*$^{um20}$ mutants, labelling all EC nuclei (green) and all arterial ECs (red). White arrowheads with numbers mark individual ECs. Note accumulation of arterialised ECs at the tip of the sprout and absence of reverse migration.

**Supplementary Movie 8. Artery-directed reverse migration of ECs is defective in *cxcr4a$^{um20}$* mutants.**

Maximum intensity projections of confocal z-stacks of *Tg(fli1a:nEGFP)$^{y7}$*; *Tg(-0.8flt1:RFP)$^{hu5333}$* double transgenic *cxcr4a$^{um20}$* mutants, labelling all EC nuclei (green) and all arterial ECs (red). White arrowheads with numbers mark individual ECs. Note accumulation of arterialised ECs at the tip of the sprout and absence of reverse migration.

**Supplementary Movie 9. Notch signalling pathway activation in artery-fated ECs.**

Maximum intensity projections of confocal z-stacks of *Tg(fli1a:nEGFP)$^{y7}$*; *Tg(-0.8flt1:RFP)$^{hu5333}$*; *Tg(TP1:Venus-Pest)$^{s940}$* triple transgenic fish labelling all EC nuclei (blue), arterial ECs (red) and Notch positive cells (green) in lateral views anterior to the left. Time-lapse imaging of the formation of the posterior circulatory loop. Note activation of Notch signalling in arterial-fated ECs (white arrowheads). Scale bar: 30 μm.

**Supplementary Movie 10. *Cxcr4a* expression in arterial-fated ECs.**

Maximum intensity projections of confocal z-stacks of *Tg(fli1a:nEGFP)$^{y7}$*; *Tg(cxcr4a:YFP)$^{mu104}$*; *Tg(-0.8flt1:RFP)$^{hu5333}$* triple transgenic fish labelling all EC nuclei (blue), arterial ECs (red) and *cxcr4a* positive cells (green) in lateral views anterior to the left. Time-lapse imaging of the formation of posterior circulatory loop. Note the expression of *cxr4a* in arterial-fated ECs migrating towards the dorsal aorta (white arrowheads). Scale bar: 20 μm.

**Supplementary Movie 11. *Cxcr4a* positive ECs display Notch pathway activation.**

Maximum intensity projections of confocal z-stacks of *Tg(fli1a:nEGFP)$^{y7}$*; *Tg(cxcr4a:YFP)$^{mu104}$*; *Tg(TP1:H2B-mcherry)$^{s939}$* triple transgenic fish labelling all EC nuclei (blue), *cxcr4a* positive cells (green) and cells with Notch pathway activation (red) in lateral views anterior to the left. Time-lapse imaging of the formation of posterior circulatory loop. Note activation of Notch signalling and *cxcr4a* in ECs migrating towards the dorsal aorta (white arrowheads). Scale bar: 30 μm.

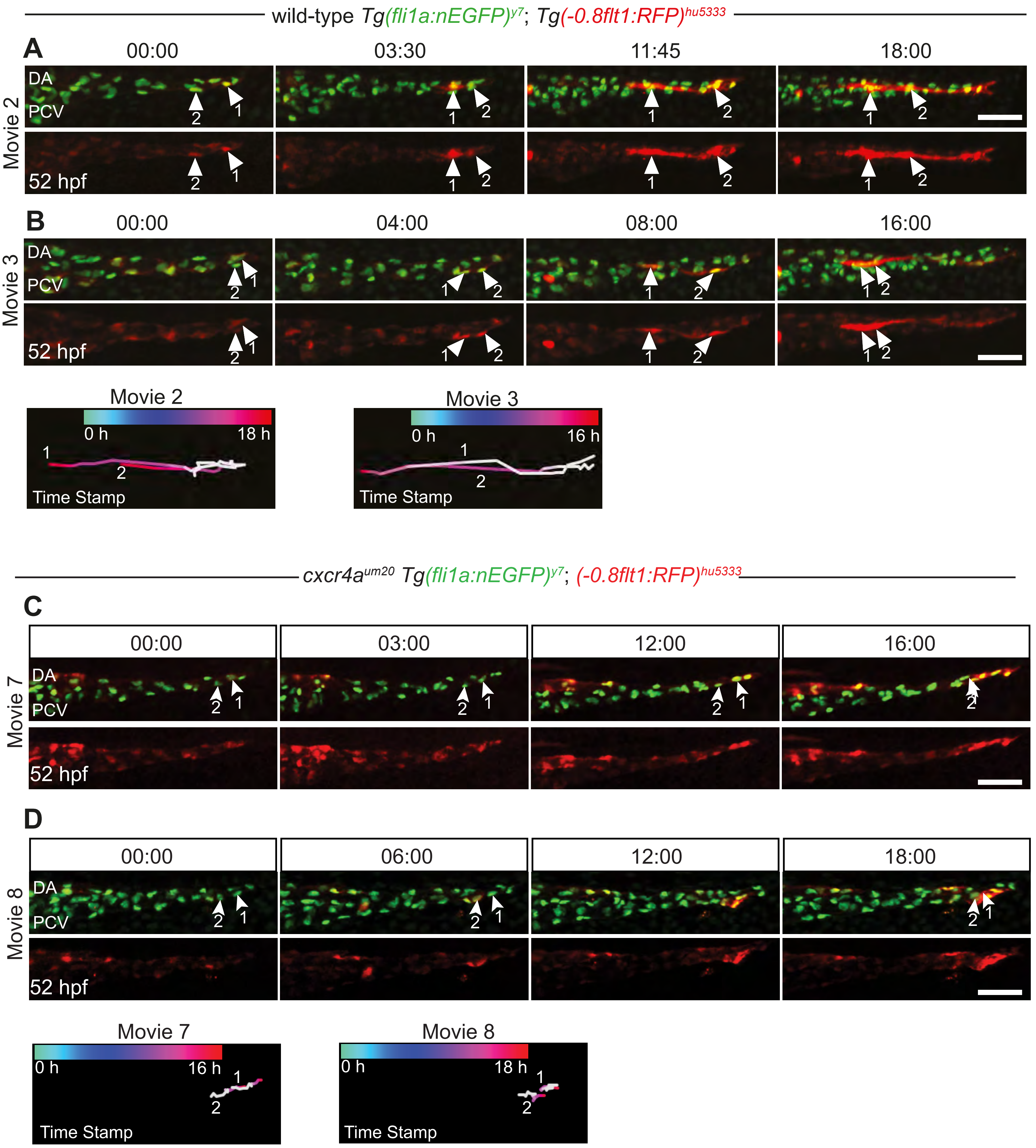


Leonard et al., Supplementary Fig.1

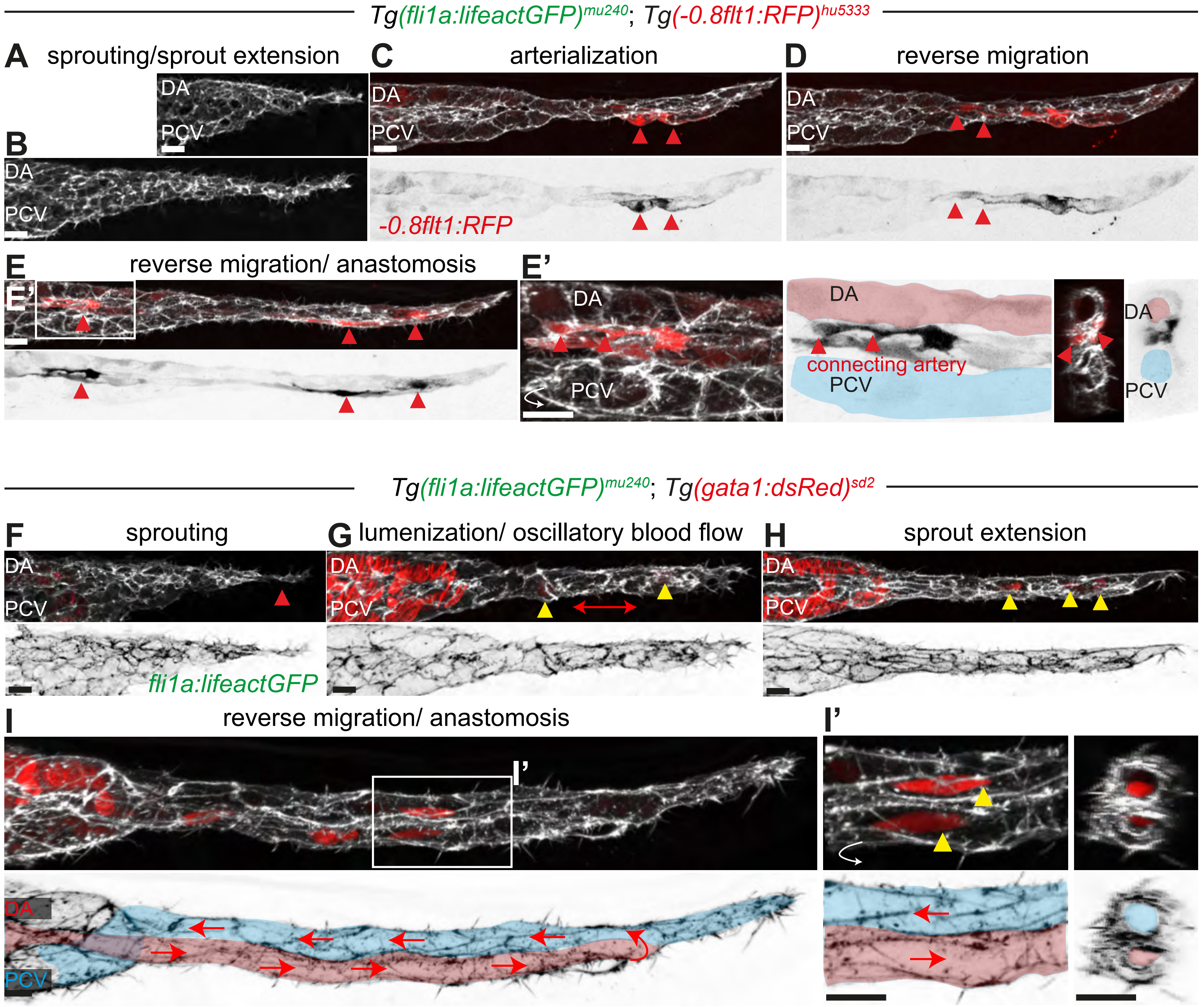


Leonard et al., Supplementary Fig. 2

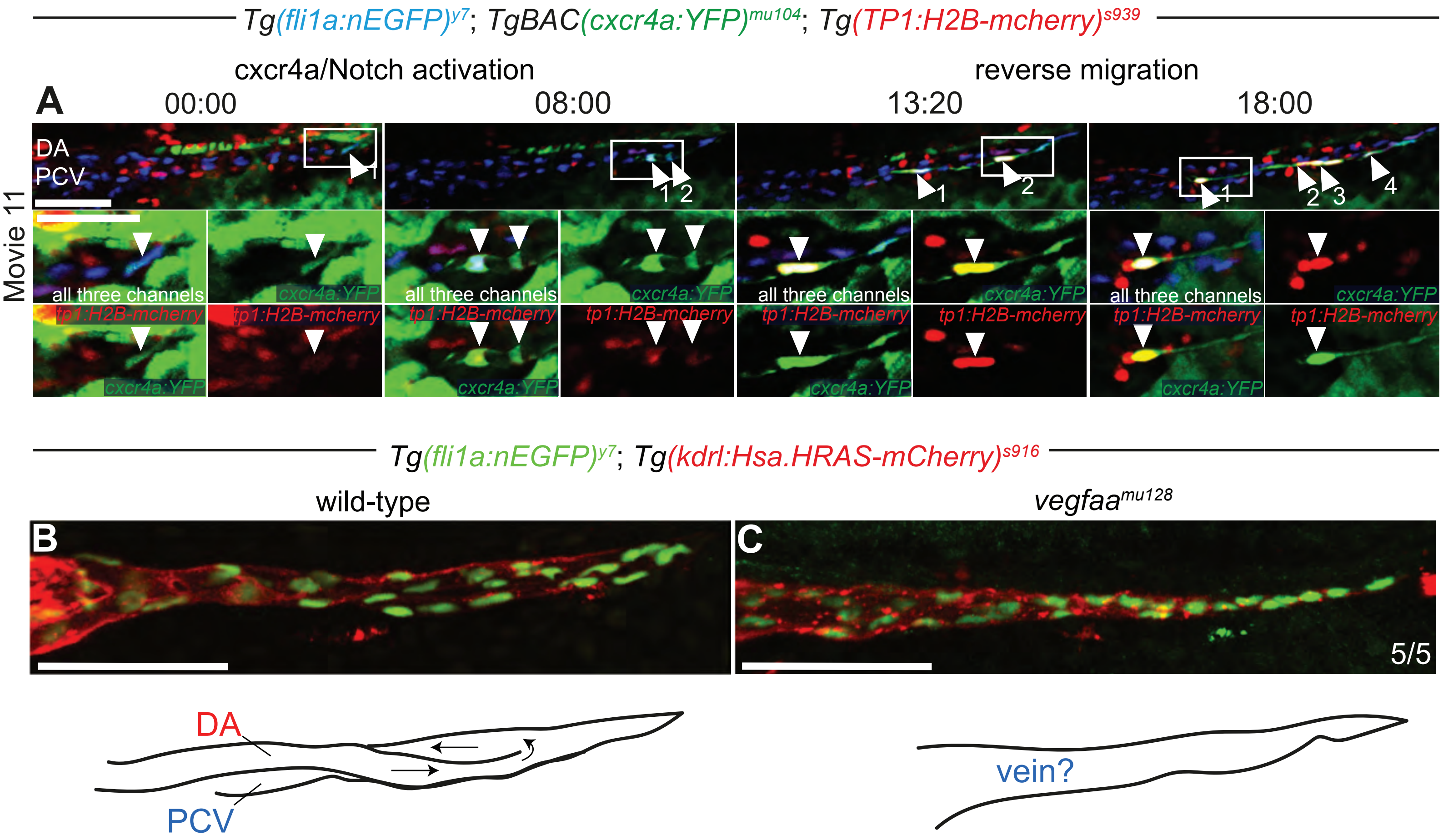


Leonard et al., Supplementary Fig. 3

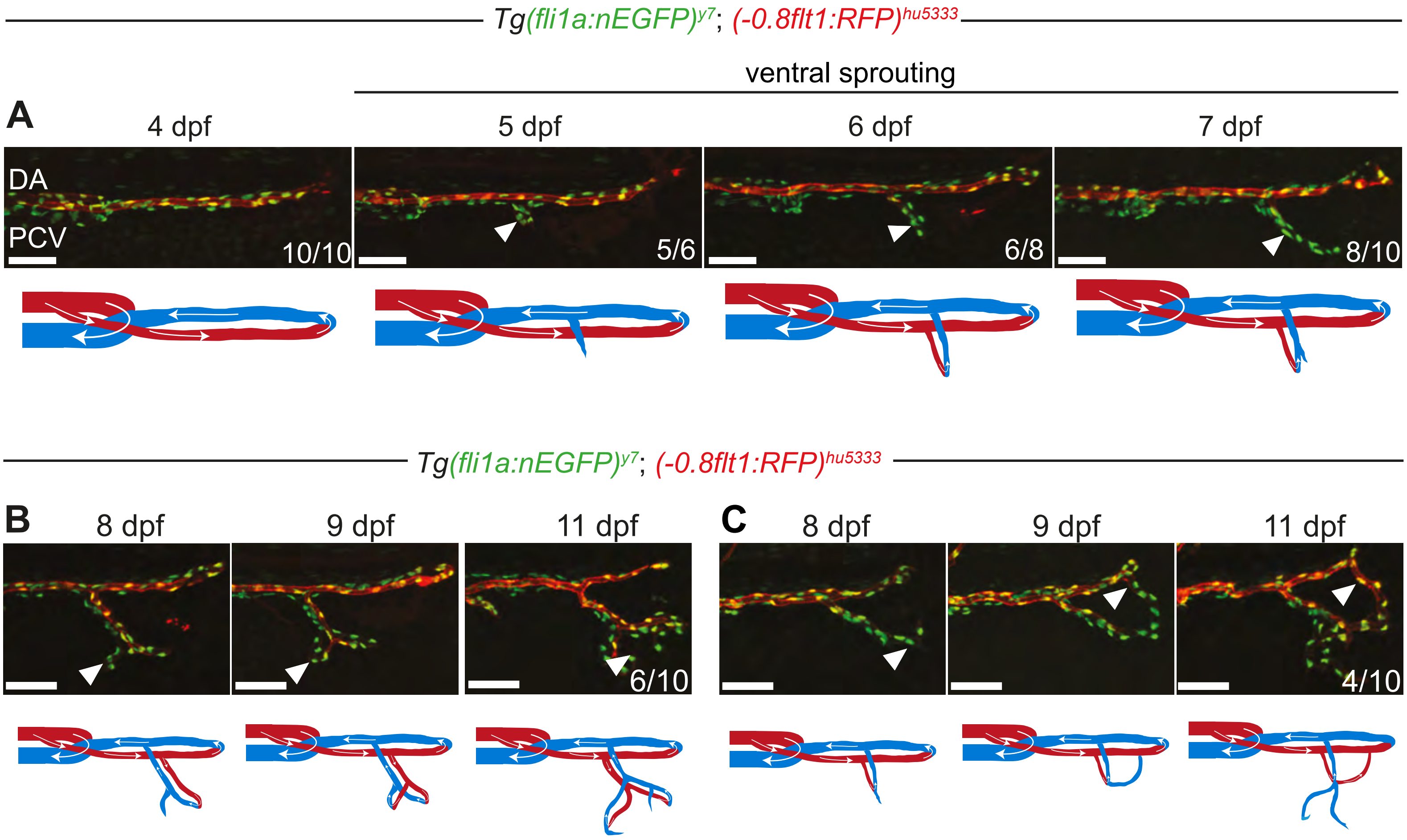


Leonard et al., Supplementary Fig.4

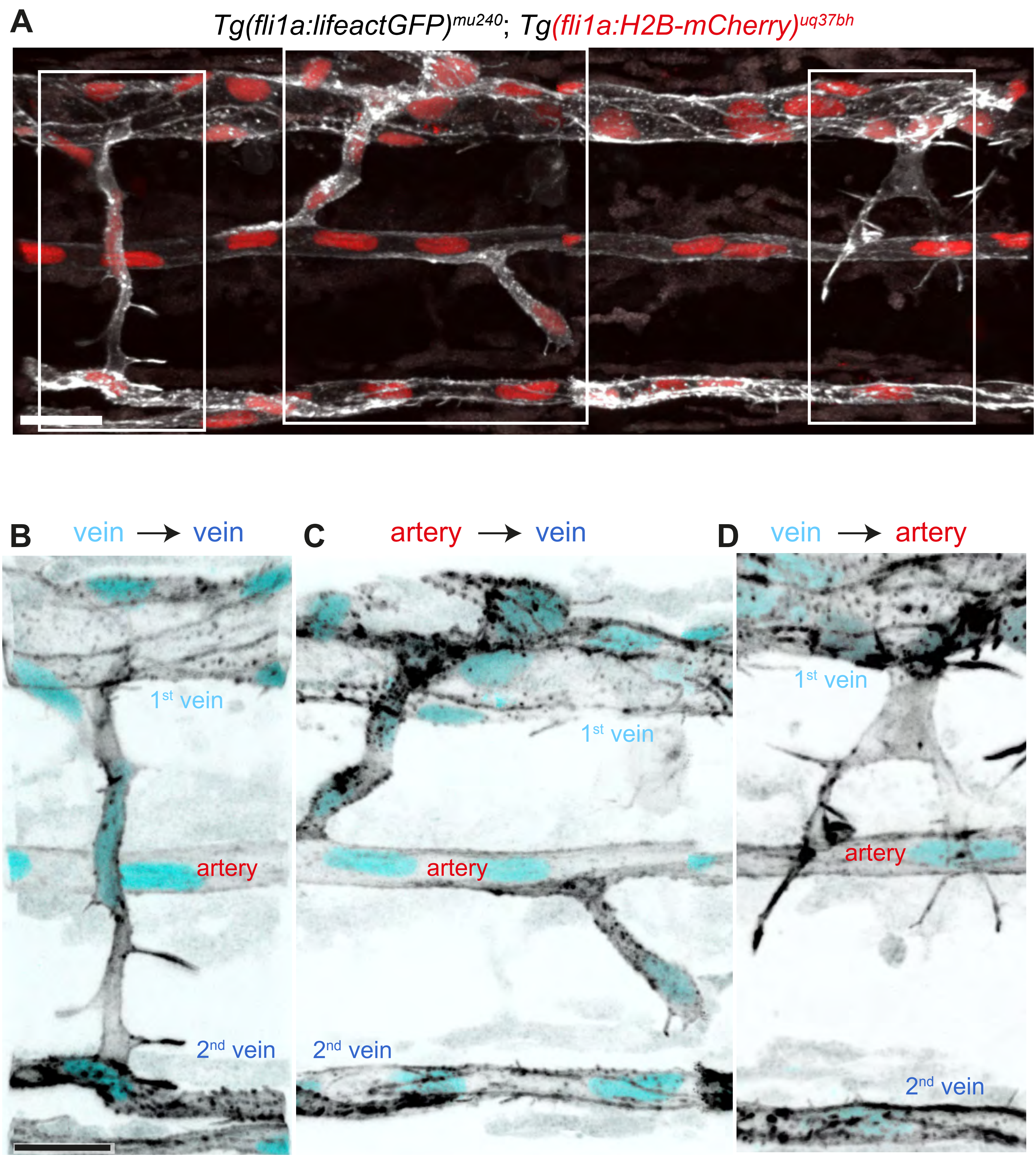


Leonard et al., Supplementary Fig.5
